# Temporal gene regulation enables controlled expression of gas vesicles and preserves bacterial viability

**DOI:** 10.1101/2025.06.20.660610

**Authors:** Zongru Li, Sumin Jeong, George J. Lu

**Affiliations:** Department of Bioengineering, Rice University, Houston, TX 77005, USA

## Abstract

Gas vesicles (GVs) are genetically encodable, air-filled protein nanostructures that have rapidly emerged as a versatile platform for biomedical imaging, cell tracking, and therapeutic delivery. However, their heterologous expression in non-native hosts such as *Escherichia coli* can be challenging due to the complex assembly process, which often involves around ten different proteins and can lead to proteotoxic stress and impair cell growth. Here, we report the observation of a drop in cell density occurring 8 to 16 hours after GV induction in *E. coli*. To address these, we developed a dual-inducer transcriptional regulation system that enables orthogonal control of GV assembly factor proteins and the shell protein over a range of stoichiometries. Sequential induction in time, in which assembly factor expression is initiated before shell protein expression, restored normal bacterial growth and prevented lysis without compromising GV production. Further analysis revealed that varying the interval between the two induction steps affected both GV yields and cellular stress. By preserving cell integrity with GV expression, our approach enhances the utility of GV-expressing bacteria in applications that demand population-wide cellular stability and facilitates their broader application in biomedical engineering and synthetic biology.

## Introduction

Synthetic biology is driving advances in biomedical innovation by harnessing engineered microorganisms as programmable platforms for diagnostics and therapeutics. From tumor-targeting probiotics^1, 2^ to CRISPR-enabled biosensors^3, 4^ and optogenetic systems^5, 6^, the field is rapidly advancing toward *in vivo* applications where intact, metabolically active cells serve as real-time reporters^7, 8^ or delivery vehicles^9^. Central to these advances is the integration of genetic circuits, such as stress-responsive elements^10^ and feedback control systems^11, 12^, that can reduce metabolic burden and optimize gene expression while preserving host cell viability and function^13^. Such cell-viability-optimized, genetically encodable modules hold the key to enabling broader use of engineered cells for real-time diagnostics^14^, targeted drug delivery^15^, and adaptive responses *in situ*^16^.

Among the genetically encodable modules rapidly advancing in recent years, gas vesicles (GVs), often known as acoustic reporter genes (ARGs)^17, 18^, are protein-based, air-filled nanostructures naturally produced inside aquatic microorganisms, including cyanobacteria and haloarchaea, for buoyancy regulation in their native environments^19, 20^. Their unique physical properties, compared to the surrounding aqueous environment of the cell, include the low density of air^20, 21^, and a large acoustic impedance mismatch^18, 22^, which have made GVs attractive for many biomedical applications. They have been extensively repurposed as genetically encoded reporters for ultrasound, magnetic resonance imaging (MRI), and optical imaging^17, 18, 22, 23^. They have also been engineered for deep-tissue acoustic cellular control^24-26^, pressure sensing^27^, and targeted delivery to lymph nodes^28^ and solid tumors^29, 30^. Additionally, GVs have been engineered into acoustic biosensors capable of detecting enzymatic activities and measuring calcium level within live cells^31, 32^ to provide insights into dynamic cellular processes. Many of these emerging technologies require GVs to be expressed and function within intact, viable cells to enable real-time, non-invasive monitoring or modulation of cellular behavior *in vivo.* As such, optimizing GV expression for live-cell contexts has become a central focus in advancing its biomedical utility.

However, producing large quantities of GVs in non-native host cells while maintaining optimal cellular physiology remains challenging due to difficulties in assembling the major shell protein, GvpA, into mature GV nanostructures. GvpA is the primary structural component of these GV nanostructures^19, 20, 33, 34^, conferring a 2–3 nm thick rigid protein shell with a highly hydrophobic inner surface that prevents water condensation^33, 34^. This shell architecture renders GvpA an amphipathic protein with poor solubility in aqueous environments. Expressing such amphipathic proteins in non-native hosts often leads to misfolding and aggregation, ultimately impairing cell growth and, in some cases, causing lysis. Accordingly, GV operons are consistently observed to encode approximately ten additional proteins, collectively known as assembly factor proteins, which mitigate cellular stress and facilitate the proper assembly of GvpA into mature GV nanostructures through a complex, multi-stage process^35^. We hypothesize that when expressing GVs in non-native hosts, the stoichiometric ratios and timing of expression between the shell protein and these assembly factor proteins are suboptimal, as the transcriptional regulation mechanisms present in native hosts may be compromised or lose functionality upon transfer to a new cellular context. This may explain why heterologous expression in *E. coli*, even under the strong T7 promoter and IPTG induction, typically yields multiple-fold lower GV production compared to native GV-producing cyanobacteria^17^. To address these challenges associated with heterologous GV expression in *E. coli*, we hypothesize that genetic circuit designs from synthetic biology can be used to mitigate cytotoxic effects and improve overall cellular viability. In this study, we first report the observation of a drop in cell density occurring 8-16 hours after GV induction in *E. coli*, accompanied by transmission electron microscopy (TEM) evidence of cell lysis in a subset of the population. Next, to address these challenges, we developed a dual-inducer transcriptional regulation system that enables independent and orthogonal control of GV assembly factor proteins and the shell protein GvpA2 in *E. coli*. This system was designed to allow precise temporal regulation of gene expression while minimizing basal transcription and avoiding unintended regulatory crosstalk. The regulatory properties of the system were validated using fluorescent reporter assays, which demonstrated tight control with low basal expression and effective orthogonality between the two inducible promoters. Leveraging this system, we established a sequential induction strategy in which assembly factors are expressed before GvpA2. This approach restored normal bacterial growth and prevented cell lysis with GV production comparable to its wild-type counterpart. Furthermore, temporal induction analysis revealed that the interval between assembly factors and shell protein expression influences GV yields and cellular stress levels. By maintaining cell viability throughout GV synthesis, this system provides a strategy for the practical application of GV-expressing bacteria in vivo, where intact and functional cells are required. These findings collectively establish a modular and tunable genetic platform for advancing GV-related technology in biomedical engineering and synthetic biology.

## Results

### *E. coli* growth inhibition during GV expression

In this work, we focused on GVs encoded by the pNL29 operon, originally cloned from a truncated portion of the GV operon in *Priestia megaterium* (formerly known as *Bacillus megaterium*)^36^. The plasmid pST39-pNL29 (**Fig. 1a**), which carries this operon, has been shown to enable efficient heterologous expression of GVs in *E. coli*^36, 37^, and has served as the workhorse for constructing acoustic reporter genes (ARGs) and determining cryo-EM structures of GVs^17, 33^. The pST39-pNL29 plasmid comprises a series of genes for GV assembly, including *gvpA2*, which encodes the primary shell protein GvpA2, followed by genes (*gvpR, N, F, G, L, S, K, J, T, U*) encoding assembly factor proteins. These genes were usually organized under the control of a T7 promoter (T7) and terminator (T7term) to enable robust GV expression in *E. coli*, which can be characterized by transmission electron microscopy (TEM) as hollow and cylindrical nanostructures after purification (**Fig. 1b**). In such systems, GV expression is induced by adding isopropyl β-D-1-thiogalactopyranoside (IPTG), activating the lacUV5 promoter that controls the chromosomally integrated T7 RNA polymerase gene (**Fig. 1c**). T7 RNA polymerase subsequently initiates transcription of the pNL29 operon through its upstream T7 promoter, resulting in the expression of GV proteins (**Fig. 1c**). Notably, the pST39-pNL29 plasmid lacks the lacI repressor and lac operator sequences, allowing for effective induction of GV expression with relatively low IPTG concentrations of 20 µM, which has been used in multiple studies as the optimized induction condition^28, 35, 37, 38^.

**Fig. 1.**
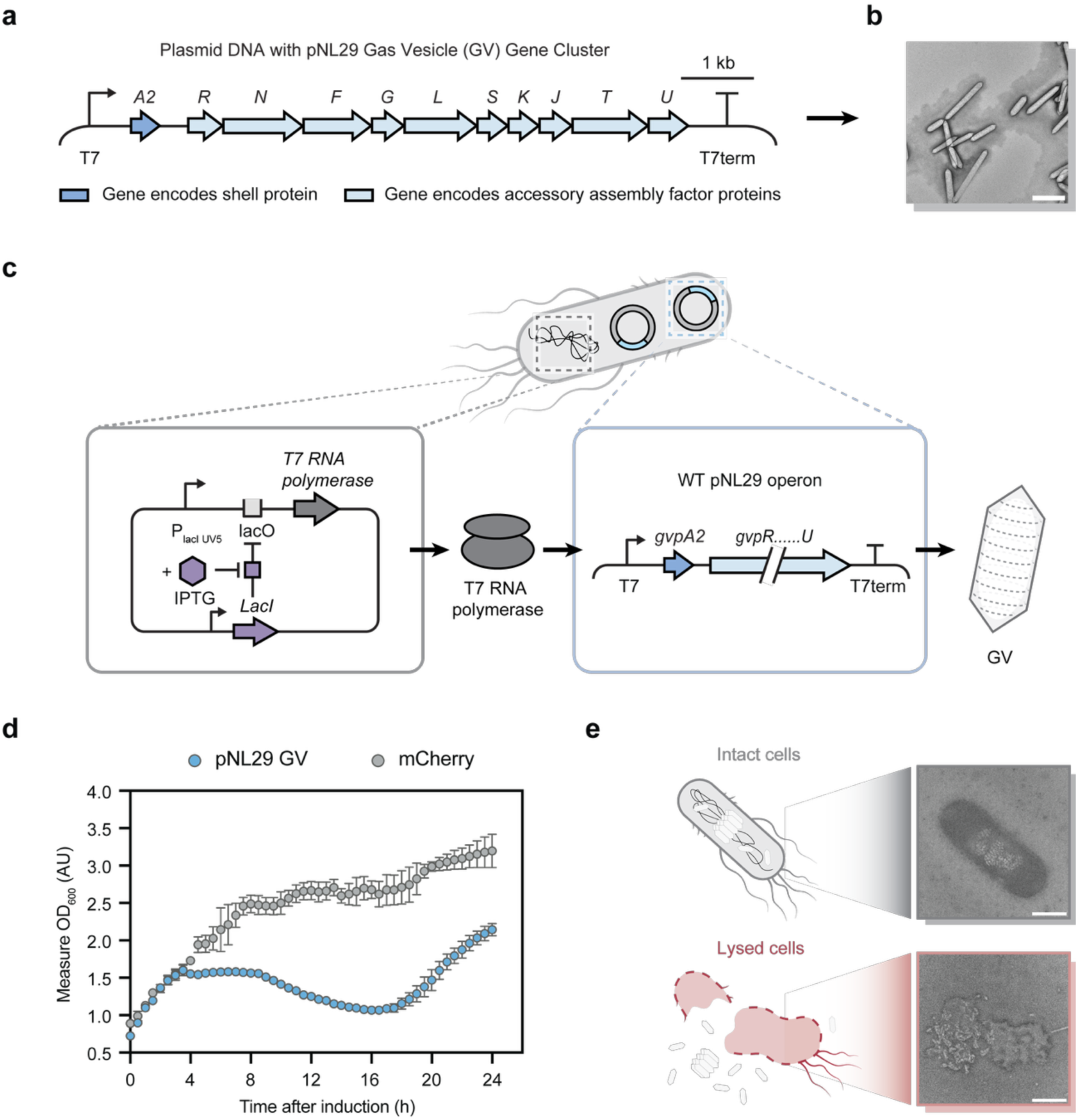
Heterologous expression of the GVs impaired bacterial growth. **a,** Schematic representation of the plasmid containing pNL29 operon architecture (pST39-pNL29). The gene encoding the shell protein GvpA2 is depicted in dark blue, while genes encoding assembly factor proteins—GvpR, GvpN, GvpF, GvpG, GvpL, GvpS, GvpK, GvpJ, GvpT, and GvpU—are shown in light blue. **b,** Representative TEM images of wild-type pNL29 GVs expressed in and purified from *E. coli*. 6 M urea treatment was applied to uncluster these GVs after purification. **c,** Schematic representation of pNL29 GVs expression in *E. coli.* Upon induction with IPTG, the lacUV5 promoter drives the expression of chromosomally integrated T7 RNA polymerase (grey box). The T7 RNA polymerase subsequently initiates transcription of the pNL29 operon under the control of a T7 promoter (blue box), leading to the expression of GVs. **d,** Growth of *E. coli* cultures after being induced with 20 µM IPTG was monitored by measuring optical density at 600 nm (Measured OD_600_). Data represent mean ± standard deviation (s.d.) for *n* = 6 biologically independent samples for the pNL29 GV-expressing culture and *n* = 4 biologically independent samples for the mCherry-expressing culture. **e,** Representative TEM images depicting an intact *E.* coli cell expressing wild-type pNL29 GVs at 4 hours post-induction (grey) and a lysed *E. coli* cell illustrating the release of GVs at 8 hours post-induction (red). Scale bars, 500 nm in **b** and **e**. Absorbance values in **d** are expressed in absorbance units (AU) on the *y-axis*, representing the amount of light absorbed by the culture at 600 nm wavelength.

However, we observed signs of compromised cellular growth in overnight cultures across multiple experimental batches. Specifically, following IPTG induction, *E. coli* cultures entered a normal exponential growth phase, but plateaued around 4 hours post-induction, followed by a marked decline in optical density commencing around 8 hours post-induction (**Fig. 1d**). This growth arrest and subsequent decline were absent in cultures without IPTG induction (**Supplementary Fig. 1**a). Moreover, increasing the IPTG concentrations to 200 and 400 µM accelerated the onset of growth arrest to approximately an hour post-induction and compounded the decline in optical density to around 2.5 hours post-induction (**Supplementary Fig. 1**b, c). In contrast, control cultures expressing the fluorescent protein mCherry under the same IPTG concentration displayed a growth trajectory consistent with typical *E. coli* growth patterns (**Fig. 1d** and **Supplementary Fig. 1**a, b, and **c**), indicating that the observed growth arrest in GV-expressing cultures is specific to GV expression and not attributable to IPTG induction.

To further confirm the hypothesis that substantial cellular toxicity exists around the 8-hour post-induction time window, we conducted TEM characterization on GV-expressing *E. coli* cells to investigate the cellular effects associated with GV expression. At 4 hours post-induction, TEM images revealed the presence of well-formed, spindle-shaped GVs within intact bacterial cells, indicating successful heterologous expression and assembly of GVs without any cytotoxic effects (**Fig. 1e**). However, by 8 hours post-induction, TEM images demonstrated significant morphological alterations, including compromised cell envelope integrity and leakage of intracellular contents, as evidenced by the presence of GVs outside of cells. These findings indicate that although the pNL29 GVs can be strongly expressed in *E. coli*, GV expression can induce a substantial physiological burden. This stress compromises cells’ structural integrity and ultimately leads to cell lysis, as demonstrated by growth inhibition and decline in optical density measurements.

### Aggregation and misfolding of GvpA2 lead to cellular proteotoxic stress

To investigate the underlying cause of proteotoxic stress observed during the expression of pNL29 GVs in *E. coli*, we focused on GvpA2, the shell protein encoded by the pNL29 operon. GvpA2 is characterized by pronounced hydrophobicity, a feature conserved among GV shell proteins^20, 39^. A recent cryo-EM study^33^ has now further elaborated that GvpA2 monomers possess an exceptionally hydrophobic interior surface that repels water (**Fig. 2a, b**). However, this hydrophobic character also underlies the tendency of GvpA2 and its homologs to aggregate and misfold when overexpressed in heterologous hosts, leading to proteotoxic stress^40^. Indeed, when expressing GvpA2 without assembly factor proteins in *E. coli*, we observed an even earlier and more significant decline in optical density (**Fig. 2c**) compared to cultures co-expressing GvpA2 with its assembly factors (**Fig. 1d**).

**Fig. 2.**
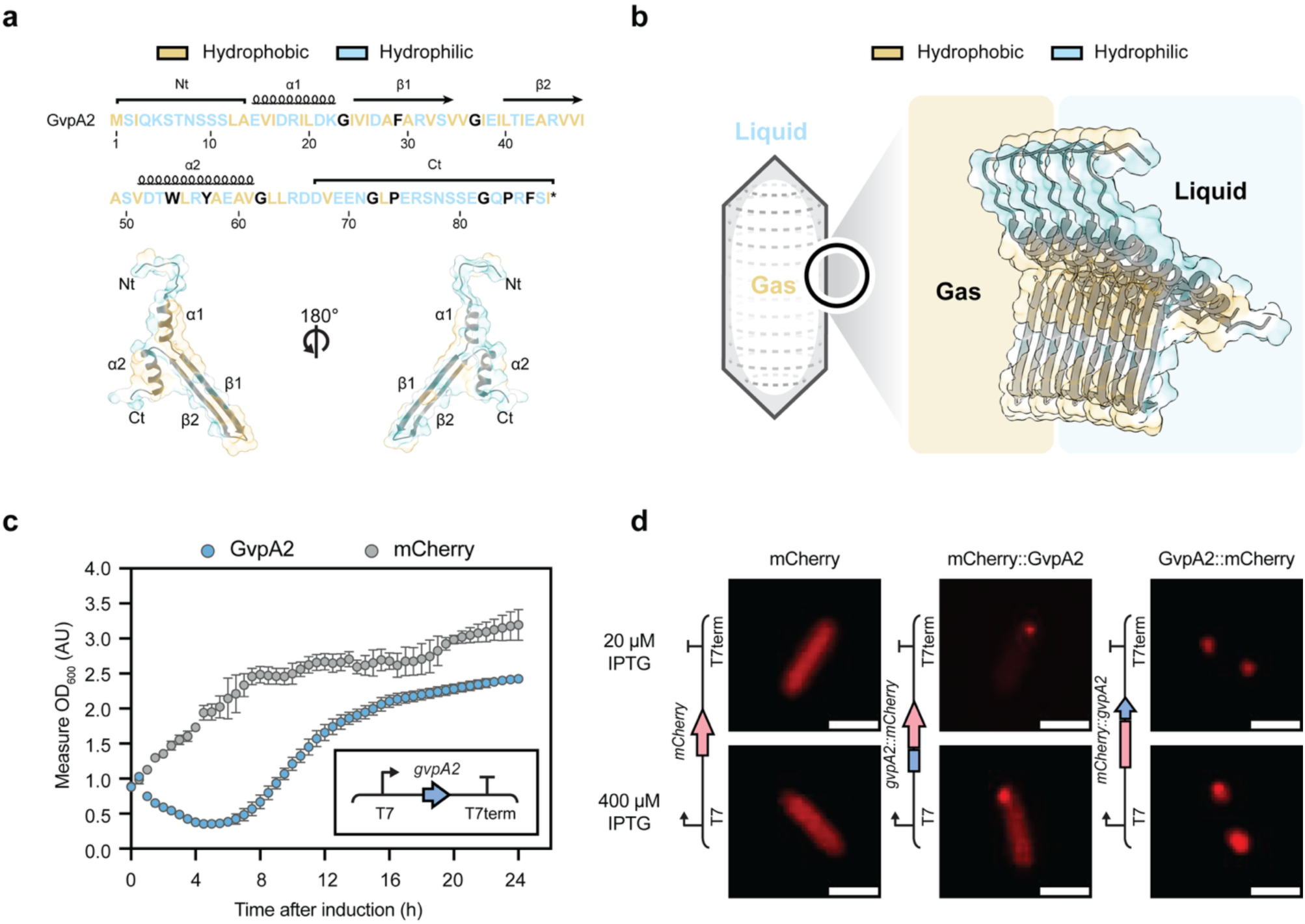
Hydrophobic shell protein GvpA2 forms aggregates in cells and *in vitro*, and triggers proteotoxic stress in heterologous expression. **a,** Amino acid sequence and cryo-EM structure^33^ (PDB 7R1C) of the major gas vesicle shell protein GvpA2. Structural fragments are annotated as N-terminal (Nt), first α-helix (α1), first β-sheet (β1), second β-sheet (β2), second α-helix (α2), and C-terminal (Ct) regions. Hydrophobic residues are depicted in yellow, while hydrophilic residues are shown in light blue. **b,** Structural organization of GvpA2 within the GV shell (PDB 7R1C). Hydrophobic amino acid residues of GvpA2 are oriented toward the GV’s interior, establishing a gas-facing surface, while hydrophilic residues are exposed on the exterior, interfacing with the aqueous environment. This amphipathic arrangement facilitates the formation of a stable gas–liquid interface, essential for GV functionality. **c,** *E. coli* growth was monitored by OD_600_ with mean ± standard deviation (s.d.) for *n* = 6 biologically independent samples for the GvpA2-expressing culture and *n* = 4 for the mCherry-expressing culture. **d,** Fluorescence microscopy images of *E. coli* cells expressing mCherry, mCherry::GvpA2, and GvpA2::mCherry. Red fluorescence images were acquired at 4 hours post-induction with 20 and 400 µM IPTG. Scale bars, 2 µm in **d**. Absorbance values in **c** are expressed in absorbance units (AU) on the *y-axis*, representing the amount of light absorbed by the culture at 600 nm wavelength.

We further set out to examine whether we can visualize the aggregates of GvpA2 in living cells^40^. To this end, we fused the mCherry fluorescent protein to either the N- or C-terminus of GvpA2 and tracked fluorescence signals through microscopy 4 hours post-induction with varying IPTG concentrations (**Fig. 2d**). Unlike the homogeneous fluorescence observed in control cells expressing mCherry alone, both mCherry::GvpA2 and GvpA2::mCherry fusions exhibited distinct fluorescent puncta (**Fig. 2d**), indicative of aggregate formation. These findings collectively indicate that the aggregation of GvpA2 can be the major cause of cellular proteotoxic stress. Addressing this challenge would therefore be crucial for optimizing the heterologous expression of GVs and mitigating their deleterious effects on host cell physiology.

### Engineering a dual-inducer system for orthogonal control of gene expression

To develop an effective strategy for alleviating this expression-associated burden, we drew inspiration from the native regulatory mechanisms that control GV expression in halophilic archaea^19, 39^. In halophilic archaea such as *Halobacterium salinarum* (*Hbt. salinarum*) and *Haloferax mediterranei* (*Hfx. mediterranei*), GV formation is intricately regulated by the interplay between two transcriptional regulators (**Fig. 3a**): GvpE, an activator, and GvpD, a repressor. These two proteins modulate the expression of gvp gene clusters in two transcriptional directions (**Fig. 3a**). GvpE binds to upstream activation sequences adjacent to gvp promoters, thereby enhancing transcription^41^. Conversely, GvpD represses GV formation by interacting with GvpE, leading to GvpE degradation and subsequent downregulation of *gvp* gene expression^41^. This GvpD and GvpE interaction also has a temporal switch: GvpD is abundant during early growth to prevent premature expression, while its decline in the late exponential phase allows GvpE accumulation and activation of *gvp* gene transcription for GV formation^42, 43^. This temporal regulatory mechanism ensures tightly controlled GV production and allows cells to respond to environmental conditions and maintain cellular homeostasis.

**Fig. 3.**
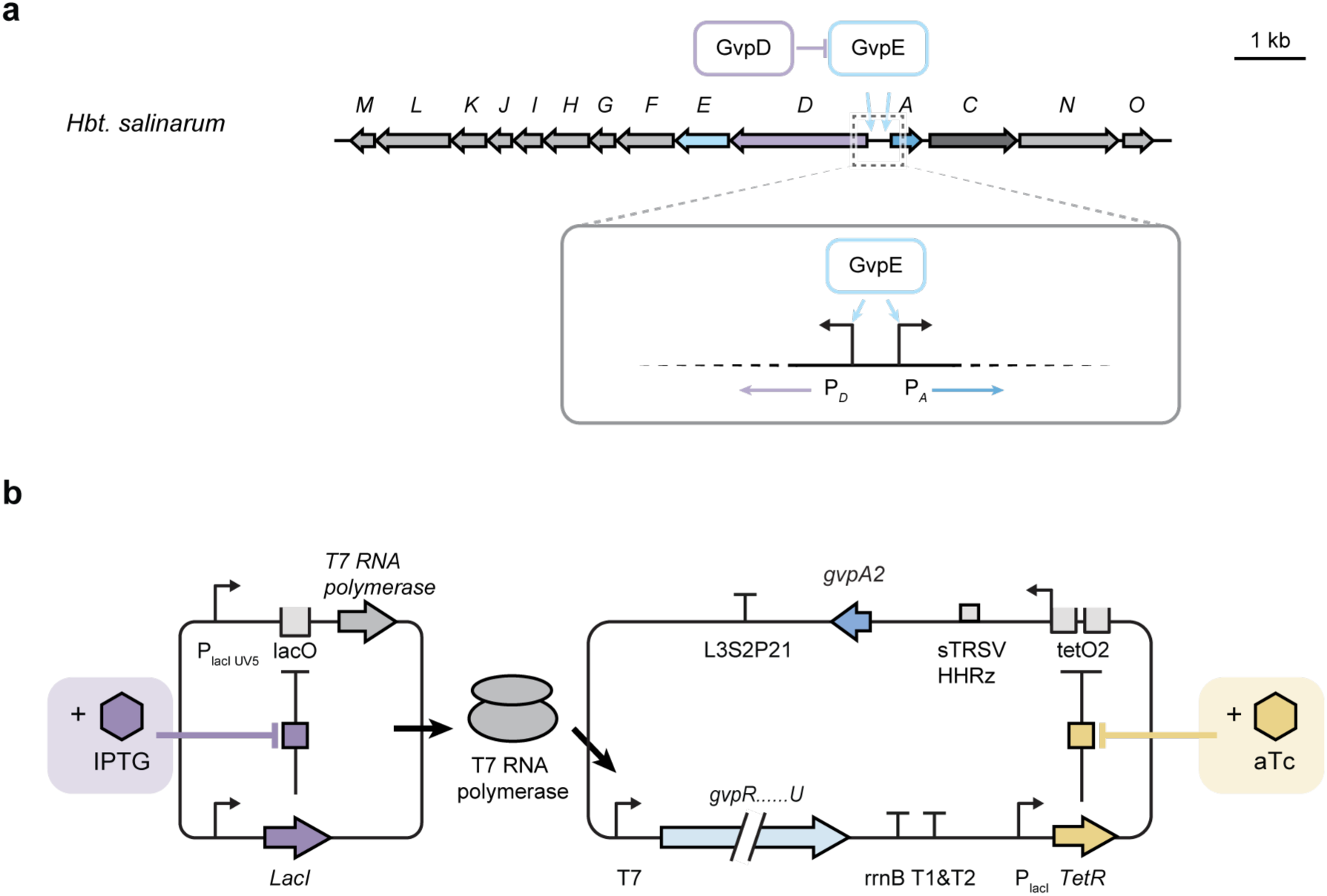
Dual-inducer transcriptional regulation enabling independent control of assembly factor and shell protein expression. **a,** Schematic representation of the gas vesicle (gvp) gene clusters in *Halobacterium salinarum* (*Hbt. salinarum*). The regulatory genes *gvpD* and *gvpE* are transcribed in opposite orientations within the cluster. GvpE (blue) functions as a transcriptional activator, enhancing the expression of structural gvp genes, while GvpD (purple) acts as a repressor by reducing the intracellular levels of GvpE, thereby modulating GV formation in *Hbt. salinarum*. **b,** Schematic representation of the dual-inducer transcriptional regulation system designed for independent control of assembly factor proteins and shell protein expression. Assembly protein genes are placed under the control of a T7 promoter, with transcription initiated by T7 RNA polymerase expression upon induction with IPTG (purple). Shell protein gene expression is regulated by a tetracycline-responsive promoter (aTc-inducible system), where transcription is repressed by the Tet repressor (TetR) in the absence of aTc and derepressed upon aTc addition (yellow).

Inspired by the native regulatory mechanisms in halophilic archaea, we developed a dual-inducer transcriptional regulation system (mentioned as ‘dual-inducer system’ in later texts; **Fig. 3b**) to mitigate the proteotoxic stress associated with heterologous GV expression in *E. coli*. In an example design, assembly factor genes can be placed under the control of a T7 promoter, with transcription initiated by T7 RNA polymerase upon induction with IPTG. Meanwhile, shell protein gene expression in the inverted transcriptional direction is regulated by a tetracycline-responsive promoter (anhydrotetracycline (aTc)-inducible system^44^), wherein transcription is repressed by the Tet repressor (TetR) in the absence of aTc and derepressed upon aTc addition. This modular system provides orthogonal induction inputs that allow for independent GV component expression dynamics (**Fig. 3b**). Specific assembly factors have been identified as potential chaperones that interact with the hydrophobic shell protein GvpA2, preventing its premature aggregation and facilitating proper assembly into the GV structure^35, 39^. These interactions suggest that the coordinated expression of assembly factors plays a crucial role in maintaining GvpA2 solubility and ensuring efficient GV expression^35^. By decoupling the expression of shell protein and assembly factors, this system aims to enable independent and temporal control over each of them, thereby minimizing the cellular proteotoxic stress from misfolded GvpA2 aggregates and improving growth inhibition during GV expression.

### Validating low basal activity and orthogonal control of the dual-inducer system by fluorescent reporters

First, we aim to establish the induction strength and orthogonality of the engineered dual-inducer system. To this end, we utilized fluorescent reporters, sfGFP and mCherry, as proxies for shell protein and assembly factors, respectively (**Fig. 4a**). Induction-specific fluorescence fold-changes were quantified for each module and compared against single-reporter controls (**Fig. 4b**). IPTG-induced mCherry expression in the dual-inducer construct exhibited lower fold-change values across all tested IPTG concentrations relative to the single-reporter control (**Fig. 4c**). This reduction likely reflects the metabolic burden^12, 45^ from constitutive TetR expression, decreasing available cellular resources. Conversely, the aTc-induced sfGFP expression in the dual-inducer system showed fold-change values comparable to the single-reporter control across all tested aTc concentrations (**Fig. 4d**), underscoring the robust independence of the aTc-inducible system regulatory module within the dual-inducer system. Next, to evaluate basal expression levels of each inducible module, we measured fluorescence intensity fold-change without induction in the dual-inducer system relative to single-reporter controls. Basal mCherry expression (0 µM IPTG; varying aTc) in the dual-inducer construct showed reduced fold-change compared to the mCherry-only control (0 µM IPTG; **Fig. 4e** and **Sup. Fig. 2**a). Similarly, basal sfGFP expression (0 ng/mL aTc; varying IPTG) closely matched the minimal baseline from the sfGFP-only control (**Fig. 4f** and **Sup. Fig. 2**b). These observations indicate minimal leakage and stringent repression of both IPTG- and aTc-inducible modules under uninduced conditions.

**Fig. 4.**
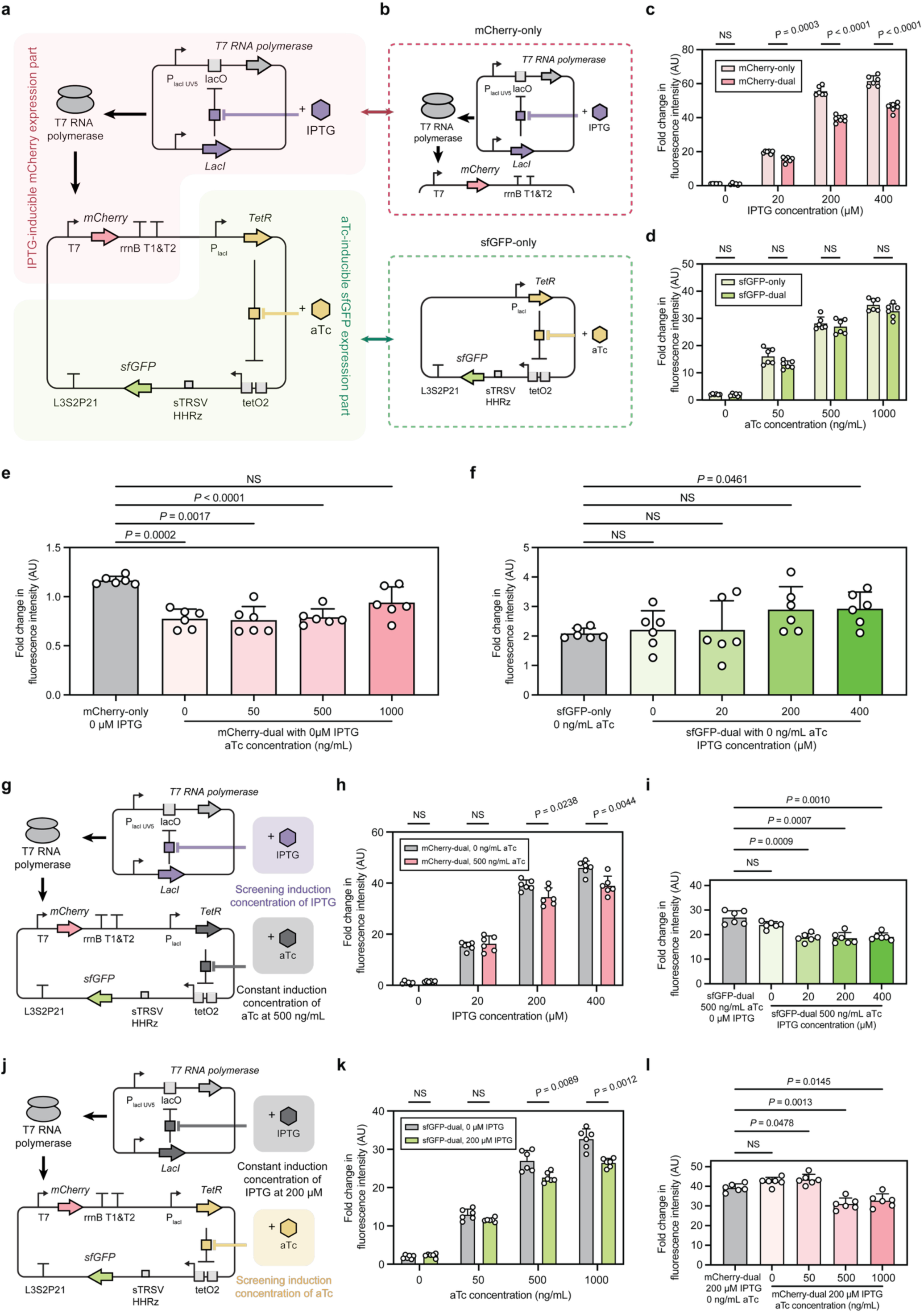
Fluorescent reporter analysis demonstrating minimal basal expression and absence of unintended regulatory interference in the dual-inducer transcriptional regulation system. **a, b,** Schematic representation of transcriptional regulation systems for expression of fluorescent reporter proteins. (**a**)The dual-inducer transcriptional regulation system enables separate control of mCherry and sfGFP expression. The mCherry expression is driven by a T7 promoter and activated upon IPTG induction (red background); sfGFP expression is regulated by an aTc-inducible system, repressed by TetR in the absence of aTc and derepressed upon aTc addition (green background). (**b**) Two single-reporter genetic constructs, with mCherry expressed upon IPTG induction (“mCherry-only,” red dashed box) and sfGFP expressed upon aTc induction (“sfGFP-only,” green dashed box), respectively. **c, d,** Fold change in fluorescence intensity comparing the dual-inducer transcriptional regulation system to single-reporter constructs (mCherry-only and sfGFP-only in **b**) under varying inducer concentrations. In the dual system, only the inducer corresponding to the respective reporter was added, resulting in “mCherry-dual” (**c**) and “sfGFP-dual” (**d**). NS, not significant (*P* > 0.05) for mCherry-only vs. mCherry-dual at 0 µM IPTG; *P* = 0.0003, < 0.0001, and < 0.0001 at 20, 200, and 400 µM IPTG, respectively. NS for all comparisons between sfGFP-only and sfGFP-dual. **e, f,** Fold change in fluorescence intensity comparing the dual-inducer transcriptional regulation system to single-reporter constructs to assess the basal expression level. (**e**) mCherry-dual groups (0 μM IPTG) with varying aTc concentrations (0, 50, 500, 1000 ng/mL) were compared to mCherry-only at 0 μM IPTG. (**f**) sfGFP-dual groups (0 ng/mL aTc) with varying IPTG concentrations (0, 20, 200, 400 μM) were compared to sfGFP-only at 0 ng/mL aTc. For mCherry-dual, *P* = 0.0002, 0.0017, and <0.0001 at 0, 50, and 500 ng/mL aTc, respectively; NS, not significant (*P* > 0.05) for 1000 ng/mL. For sfGFP-dual, *P* = 0.0461 at 400 μM IPTG; NS for all other comparisons. **g,** Schematic representation of the genetic construct for fluorescent reporter proteins under the dual-inducer transcriptional regulation system. Cultures were induced with varying concentrations of IPTG (0, 20, 200, and 400 μM) while maintaining a constant aTc concentration of 500 ng/mL. **h,** Fold change in mCherry fluorescence intensity in the dual-inducer transcriptional system with and without aTc. mCherry-dual groups with 0 ng/mL aTc were compared to groups with 500 ng/mL aTc across varying IPTG concentrations (**g**). NS for comparisons at 0 and 20 µM IPTG; *P* = 0.0238 and 0.0044 for comparisons at 200 and 400 µM IPTG. **i,** Fold change in sfGFP fluorescence intensity in the dual-inducer transcriptional system with and without IPTG. sfGFP-dual groups with 500 ng/mL aTc and 0 µM IPTG were compared to those with 500 ng/mL aTc across varying IPTG concentrations. NS for comparisons at 0 µM IPTG; *P* = 0.0009, 0.0007, and 0.0010 for comparisons at 20, 200, and 400 µM IPTG. **j,** Schematic representation of the genetic construct for fluorescent reporter proteins under the dual-inducer transcriptional regulation system. Cultures were induced with varying concentrations of aTc (0, 50, 500, and 1000 ng/mL) while maintaining a constant IPTG concentration of 200 µM. **k,** Fold change in sfGFP fluorescence intensity in the dual-inducer transcriptional system with and without aTc. sfGFP-dual groups with 0 µM IPTG were compared to groups with 200 µM IPTG across varying aTc concentrations (**j**). NS for comparisons at 0 and 50 ng/mL aTc; *P* = 0.0089 and 0.0012 for comparisons at 500 and 1000 ng/mL aTc. **l,** Fold change in mCherry fluorescence intensity in the dual-inducer transcriptional system with and without IPTG. mCherry-dual groups with 200 µM IPTG and 0 ng/mL aTc were compared to those with 200 µM IPTG across varying aTc concentrations. NS for comparisons at 0 ng/mL aTc; *P* = 0.0478, 0.0013, and 0.0145 for comparisons at 50, 500, and 1000 ng/mL aTc. Fold change in fluorescence intensity values in **c**, **d, e**, **f, h**, **i, k,** and **l** are expressed in arbitrary units (AU) on the *y-axis*. Data in **c**, **d, e**, **f, h**, **i, k,** and **l** represent mean ± s.d. for *n* = 6 biologically independent samples for all groups. The statistical significance between groups in **c**, **d, h**, and **k** was tested using a two-tailed unpaired Student’s *t*-test with Welch correction. The statistical significance was tested against control groups (grey) in **e**, **f**, **i,** and **l** using Brown–Forsythe and Welch one-way ANOVA tests, followed by Dunnett T3 multiple comparisons correction. *P* values of each comparison are indicated in each caption.

We next assessed potential regulatory interference within the dual-inducer system under simultaneous induction conditions. At constant aTc induction (500 ng/mL), fold-change in mCherry fluorescence remained comparable between cultures with and without aTc at lower IPTG levels (0 and 20 µM, **Fig. 4g, h**). However, higher IPTG concentrations (200 and 400 µM) exhibited reduced mCherry fluorescence upon aTc co-induction (**Fig. 4h**), indicative of resource competition. A similar trend in inducibility and resource competition was observed for sfGFP (**Fig. 4i**) and in the other half of the dual-inducer system (**Fig. 4j–l**). To further preclude potential biases arising from the differing fluorescence properties of sfGFP and mCherry, we interchanged their regulatory controls within the dual-inducer system (**Sup. Fig. 3**a). In this configuration, sfGFP expression is driven by the T7 promoter and induced by IPTG, while mCherry expression is regulated by the aTc-inducible system. Results were consistent with previous observations on functionality, basal expression levels, and unintended regulatory interference (**Sup. Fig. 3** and 4). Collectively, these findings validate that the dual-inducer system maintains precise regulatory control, minimal basal expression, and minimal unintended interference, as demonstrated by its consistent performance irrespective of the specific fluorescent reporters employed.

### Expressing assembly factors before the shell protein alleviates proteotoxic stress and restores *E. coli* growth

After validating the regulatory performance of the engineered dual-inducer system, we aimed to assess whether implementing this system to achieve sequential expression of assembly factor proteins before shell protein synthesis could alleviate proteotoxic stress and restore *E. coli* growth during GV expression. Based on the rapid protein accumulation typically observed within 2–4 hours in T7-driven systems^46^, we selected a 3-hour interval between each induction to ensure sufficient expression of assembly factors to effectively chaperone GvpA2^35^. To further investigate how varying the expression levels of assembly factor proteins and shell protein influence *E. coli* growth, we adopted an analogous inducer concentration screening strategy employed in the previous fluorescent reporter analyses (**Fig. 4g, j**). Specifically, cultures were initially induced with a range of IPTG concentrations, followed by a constant aTc concentration of 500 ng/mL after 3 hours (**Fig. 5a**). In another set of screening, cultures were first induced with a fixed IPTG concentration of 200 µM, and subsequently exposed to varying aTc concentrations after the same 3-hour interval (**Fig. 5b**). For the IPTG-regulated assembly factors induction screening, all groups except the uninduced one (0 µM IPTG) exhibited restored *E. coli* growth patterns, characterized by the absence of the growth arrest or optical density decline observed in cells expressing the wild-type pNL29 GV operon (**Fig. 5c**). In contrast, the 0 µM IPTG exhibited a marked decline in optical density from approximately 5 hours after aTc-regulated induction of the shell protein to the second day. Notably, this decline occurred later than in cells expressing GvpA2 alone under the T7-driven system (**Fig. 2b**). One plausible explanation to this observation is that the aTc-inducible system, which utilizes endogenous *E. coli* RNA polymerase^47^, exhibits comparatively lower expression levels than the T7-driven system, as T7 RNA polymerase transcribes roughly 5–8 times faster than *E. coli* RNA polymerase^46, 48^. For the aTc-regulated assembly factors induction screening, all experimental groups exhibited restored *E. coli* growth profiles, with only a minimal decrease in optical density observed approximately 2 hours after aTc-induced shell protein expression (**Fig. 5d**). Meanwhile, the optical density values for both screenings at the 24-hour post-induction harvest point were consistently lower than those observed in the mCherry control group (**Fig. 1d**). This observation suggests that the expression of complex nanostructures, such as GVs, imposes a metabolic burden on *E. coli*^49^, potentially limiting maximal cell density. Nevertheless, these results demonstrate that sequential induction of assembly factor proteins before shell proteins using the dual-inducer system effectively alleviates proteotoxic stress and restores bacterial growth during GV expression.

**Fig. 5.**
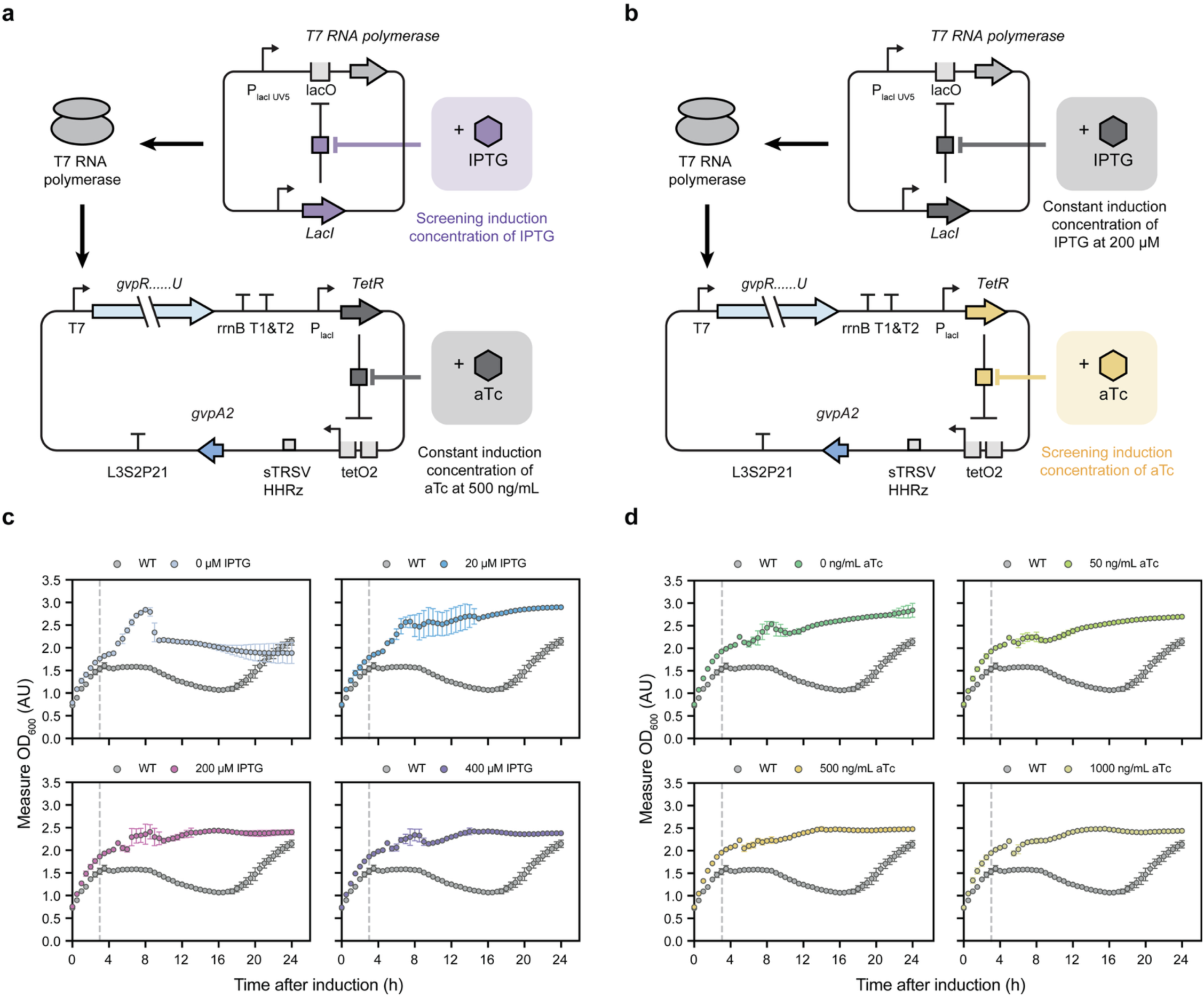
Sequential expression of assembly factor proteins prior to shell protein synthesis mitigates proteotoxic stress and restores normal growth in *E. coli*. **a,** Schematic representation of the genetic construct for GV expression under the dual-inducer transcriptional regulation system. Cultures were induced with varying concentrations of IPTG (0, 20, 200, and 400 μM) while maintaining a constant aTc concentration of 500 ng/mL. IPTG was added when the OD_600_ reached 0.6–0.8, followed by the addition of aTc three hours later. **b,** Schematic representation of the genetic construct for GV expression under the dual-inducer transcriptional regulation system. Cultures were induced with varying concentrations of aTc (0, 50, 500, and 1000 ng/mL) while maintaining a constant IPTG concentration of 200 µM. IPTG was added when the OD_600_ reached 0.6–0.8, followed by the addition of aTc three hours later. **c,** Growth profiles (measured OD_600_) of *E. coli* cultures expressing GVs under the dual-inducer transcriptional regulation system after being induced with varying IPTG concentrations and constant concentration of aTc at 500 ng/mL. The growth profile of wild-type pNL29 GV-operon-expressing *E. coli* cultures induced with 20 μM IPTG (WT) served as the control. The vertical grey dashed line indicates the time point (three hours after IPTG addition) at which aTc was introduced. **d,** Growth profiles (measured OD_600_) of *E. coli* cultures expressing GVs under the dual-inducer transcriptional regulation system after being induced with constant IPTG concentration at 200 µM and varying concentrations of aTc. The growth profile of wild-type pNL29 GV-operon-expressing *E. coli* cultures induced with 20 μM IPTG (WT) served as the control. The vertical grey dashed line indicates the time point (three hours after IPTG addition) at which aTc was introduced. Absorbance values in **b** and **d** are expressed in absorbance units (AU) on the *y-axis*, representing the amount of light absorbed by the culture at 600 nm wavelength. Data in **b** and **d** represent mean ± s.d. for *n = 6* biologically independent samples for all groups.

### Interchanged regulatory control limits the mitigation of proteotoxic stress in *E. coli*

Building upon the demonstrated interchangeability of the dual-inducer system using fluorescent proteins, we sought to evaluate whether swapping the regulatory controls of assembly factor proteins and the shell protein would yield comparable mitigation of proteotoxic stress. In the early ARG development^17^, the entire GV operon was placed under IPTG control, which led us to hypothesize that maintaining IPTG control for the primary shell protein, GvpA2, would similarly optimize GV yield. Thus, we designed a configuration in which GvpA2 expression was induced by IPTG, while assembly factor expression was controlled by aTc (**Fig. 6a**). However, the outcome was somewhat unexpected. Optical density profiles were assessed using the same methodology as previously described, wherein the concentration of one inducer was held constant while varying the concentration of the other, with a 3-hour interval between two inductions (**Fig. 6b, d**). Although the growth arrest characteristic of the wild-type was not observed, all experimental groups, except the uninduced GvpA2 group (0 µM IPTG), displayed a noticeable decrease in optical density commencing approximately 3–9 hours post IPTG-regulated shell protein induction (**Fig. 6c, e**). This observation may also be attributed to the inherently lower transcriptional strength of the aTc-inducible system^47^ compared to the T7-driven system^46, 48^. Consequently, the limited availability of assembly factor proteins, expressed under the aTc-inducible system during the preceding 3-hour induction interval, was insufficient to effectively solubilize the rapidly accumulating GvpA2 produced from the T7-driven system. As a result, the interchanged regulatory configuration did not completely mitigate proteotoxic stress induced by GvpA2, thereby failing to fully restore *E. coli* growth during heterologous expression of GV.

**Fig. 6.**
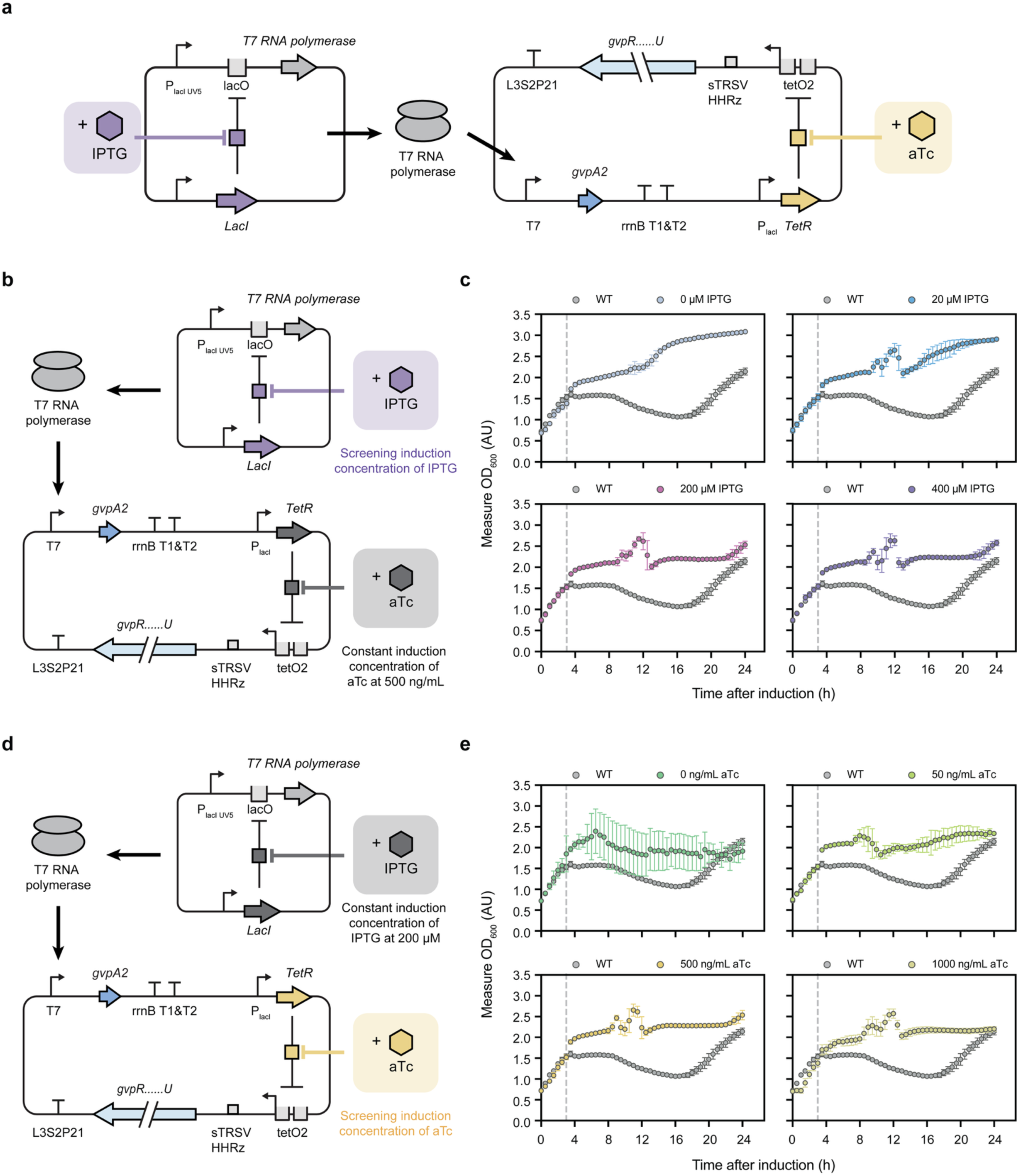
Interchanging regulatory control of assembly factor proteins and shell protein expression in the dual-inducer transcriptional regulation system resulted in limited mitigation of proteotoxic stress. **a,** Schematic representation of the interchanged dual-inducer transcriptional regulation system designed for independent control of assembly proteins and shell protein expression. Shell protein gene is placed under the control of a T7 promoter, with transcription initiated by T7 RNA polymerase expression upon induction with IPTG (purple). Assembly protein genes expression is regulated by an aTc-inducible system, where transcription is repressed by the TetR in the absence of aTc and derepressed upon aTc addition (yellow). **b,** Schematic representation of the genetic construct for GV expression under the interchanged dual-inducer transcriptional system. Cultures were induced with varying concentrations of IPTG (0, 20, 200, and 400 μM) while maintaining aTc at a constant concentration of 500 ng/mL. IPTG was added when the OD_600_ reached 0.6–0.8, followed by the addition of aTc three hours later. **c,** Measured OD_600_ of *E. coli* cultures expressing GVs under the interchanged dual-inducer transcriptional regulation system after being induced with varying IPTG concentrations and constant concentration of aTc at 500 ng/mL. The growth profile of WT 20 μM IPTG served as a control (WT). The vertical grey dashed line indicates the time point (three hours after IPTG addition) at which aTc was introduced. **d,** Schematic representation of the genetic construct for GV expression under the interchanged dual-inducer transcriptional system. Cultures were induced with varying concentrations of aTc (0, 50, 500, and 1000 ng/mL) while maintaining IPTG at a constant concentration of 200 µM. IPTG was added when the OD_600_ reached 0.6–0.8, followed by the addition of aTc three hours later. **e,** Measured OD_600_ of *E. coli* cultures expressing GVs under the interchanged dual-inducer transcriptional regulation system after being induced with constant IPTG concentration at 200 µM and varying concentrations of aTc. The growth profile of WT 20 μM IPTG served as a control (WT). The vertical grey dashed line indicates the time point (three hours after IPTG addition) at which aTc was introduced. Absorbance values in **c** and **e** are expressed in absorbance units (AU) on the *y-axis*, representing the amount of light absorbed by the culture at 600 nm wavelength. Data in **c** and **e** represent mean ± s.d for *n* = 6 biologically independent samples for all groups.

### The dual-inducer transcriptional regulation system achieves GV yields comparable to the wild-type operon

Having established the efficacy of the dual-inducer system in mitigating proteotoxic stress and restoring bacterial growth, we next sought to assess whether the sequential expression of assembly factors and shell protein influences the yield of GV production in *E. coli*. The hydrostatic collapse assay is a quantitative method used to assess GV yield by measuring the susceptibility of GVs to pressure-induced collapse^38, 50^. In this technique, a suspension of purified GVs is subjected to incrementally increasing hydrostatic pressure, and the corresponding decrease in optical density at 500 nm (OD_500_) is monitored (**Fig. 7a**). The reduction in OD_500_ reflects the collapse of GVs, as intact GVs scatter light^20^, resulting in a turbid appearance, whereas collapsed ones lose this scattering property, yielding a clear solution (**Fig. 7b**). The pressure-dependent changes in OD_500_ of GV samples can be fitted using a Boltzmann sigmoidal regression model^37^, with the goodness-of-fit assessed through the coefficient of determination (*R*²) (**Fig. 7c**). We lysed cells expressing gas vesicles (GVs) at harvest and subsequently employed the hydrostatic collapse assay to quantitatively determine GV yield in each culture, as GVs are the primary cellular component responsible for OD_500_ reduction. The hydrostatic collapse profile of cultures expressing the wild-type pNL29 GV operon induced with 20 µM IPTG served as the control, due to this group’s highest coefficient of determination (*R*² = 0.9488; **Fig. 7d-f**), indicative of substantial GV production. Interestingly, at 0 µM IPTG induction, the *R*² value (0.6012; **Fig. 7d**) was higher than that observed at 200 µM IPTG (*R*² = 0.1780; **Fig. 7e**) and 400 µM IPTG (*R*² = 0.2482; **Fig. 7f**). This observation suggests the possibility of a slight basal expression of GV proteins in the pNL29 operon, leading to limited GV formation. Meanwhile, increased IPTG concentrations intensify GvpA2-induced proteotoxic stress and cell lysis (**Sup. Fig. 1**b, c), significantly impairing GV assembly.

**Fig. 7.**
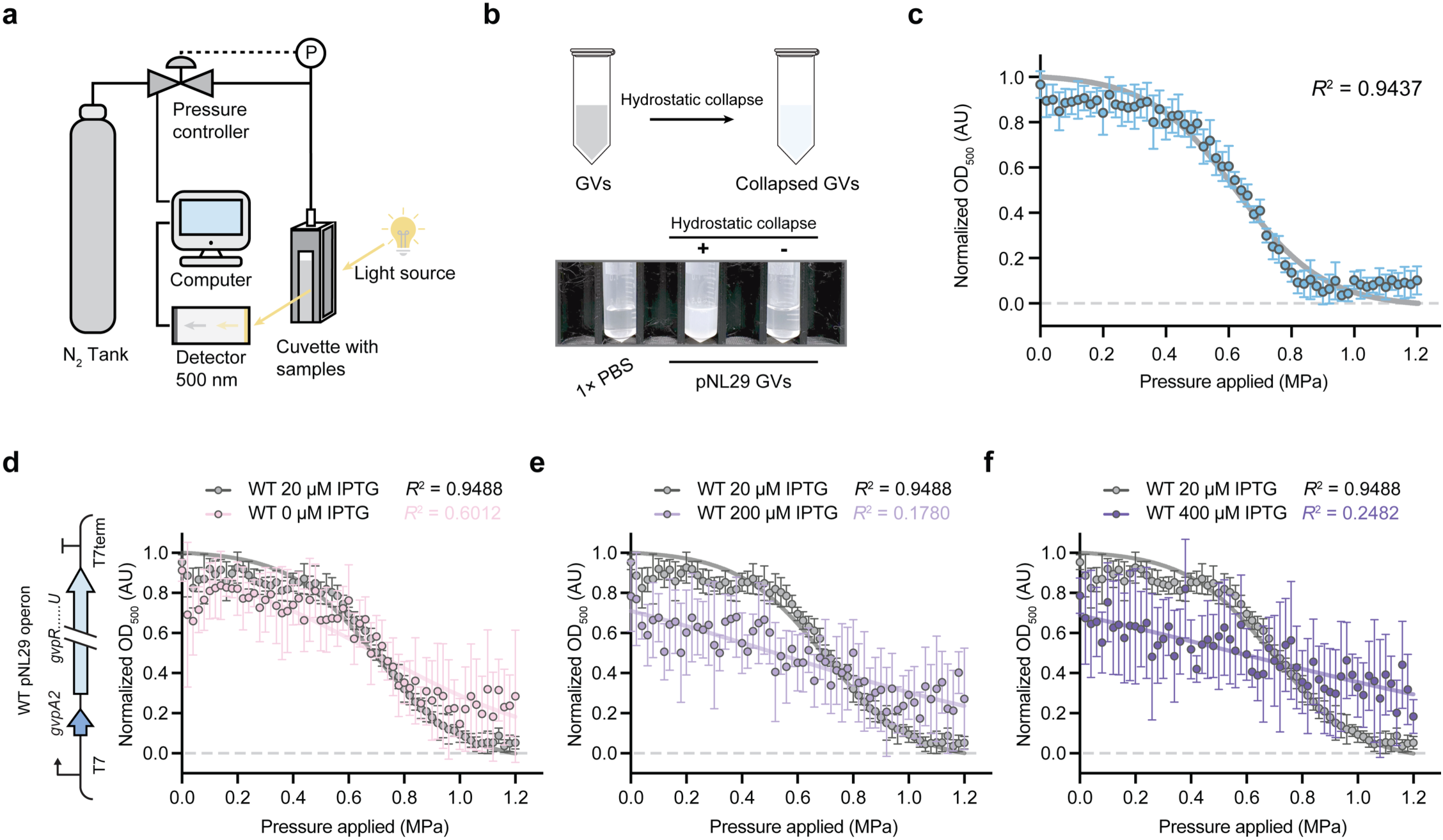
Quantitative assessment of GV yield in *E. coli* by hydrostatic collapse assay. **a,** Schematic representation of the hydrostatic collapse assay setup used to analyze GV samples. The system comprises a Nitrogen (N_2_) tank with a pressure controller connected to a sealed optical cuvette containing the suspended GV samples. Hydrostatic pressure is incrementally applied, and the optical density at 500 nm (OD_500_) is monitored using a spectrophotometer to assess the collapse behavior of GVs under increasing pressure. **b,** Schematic representation and representative images of 6 M urea-treated wild-type pNL29 GV samples before (-) and after (+) hydrostatic collapse. The GV samples exhibit decreased turbidity, corresponding to a reduced optical density value after hydrostatic collapse. **c,** Hydrostatic collapse profile for purified 6 M urea-treated wild-type pNL29 GV samples. The hydrostatic collapse data were fitted using a Boltzmann sigmoidal regression model, depicted as a grey curve, yielding a coefficient of determination (*R*²) of 0.9437. **d, e, f,** Hydrostatic collapse profiles of wild-type pNL29 GV-operon-expressing *E. coli* cultures. Cultures were induced with 0 µM (**d**), 200 µM (**e**), and 400 µM (**f**) IPTG. WT 20 µM IPTG serves as the control. Data represent mean ± s.d for *n* = 6 biologically independent samples. All data were fitted using a Boltzmann sigmoidal regression model (curves with respective colors), with corresponding *R*² values indicated for each IPTG concentration. Normalized absorbance values in **a**, **b**, and **c** are expressed in absorbance units (AU) on the *y-axis*. Normalized absorbance values in **c**, **d, e**, and **f** are expressed in absorbance units (AU) on the *y-axis*. Data in **c**, **d, e**, and **f** represent mean ± s.d for *n* = 6 biologically independent samples for all groups.

Next, to ask whether the sequential expression impacts GV production yield in *E. coli* using the dual-inducer system, we utilized the previously described protocol, maintaining a 3-hour interval between inductions while varying the concentration of one inducer and holding the other constant (**Fig. 5a, b**). In the IPTG-regulated assembly factors induction screening, all induced groups (20, 200, and 400 µM IPTG) exhibited hydrostatic collapse profiles comparable to the value of the wild-type pNL29 GV operon induced at 20 µM IPTG, as demonstrated by comparable *R*² values to the wild-type counterpart (**Fig. 8a**). Notably, the *R*² value at 20 µM IPTG (*R*² = 0.8103) is not as high as those at higher IPTG concentrations (*R*² = 0.9126 at 200 µM IPTG; *R*² = 0.9262 at 400 µM IPTG). This observation indicates a lower GV yield at 20 µM IPTG, consistent with the previously recorded optical density profiles showing higher cell densities at this induction level (20 µM IPTG, **Fig. 5c**), which reflects a reduced metabolic burden due to lower GV expression. To further substantiate these results, we calculated the reduction in OD_500_ and normalized it to OD_600_ at harvest to account for variations in cell density. As higher normalized reduction values correlated with increased GV yield, the results (**Fig. 8b**) demonstrated consistent trends with the hydrostatic collapse profiles. All IPTG-induced groups exhibited normalized reduction values comparable to the wild-type, demonstrating the efficacy of the dual-inducer system in maintaining GV production. In the aTc-regulated shell protein induction screening, groups induced at higher aTc concentrations (500 ng/mL, *R*² = 0.9303; 1000 ng/mL, *R*² = 0.9431) exhibited hydrostatic collapse profiles comparable to that of the wild-type control (**Fig. 8c**). In contrast, the low R² value observed at 50 ng/mL aTc (*R*² = 0.3162) indicates a marginal GV yield, further supported by the lower normalized reduction in OD_500_ compared to groups induced with higher aTc concentrations (**Fig. 8d**). This reduced yield is likely due to the limited availability of the GvpA2 expressed under the aTc-inducible system at lower aTc concentrations, resulting in insufficient shell protein for efficient GV assembly.

**Fig. 8.**
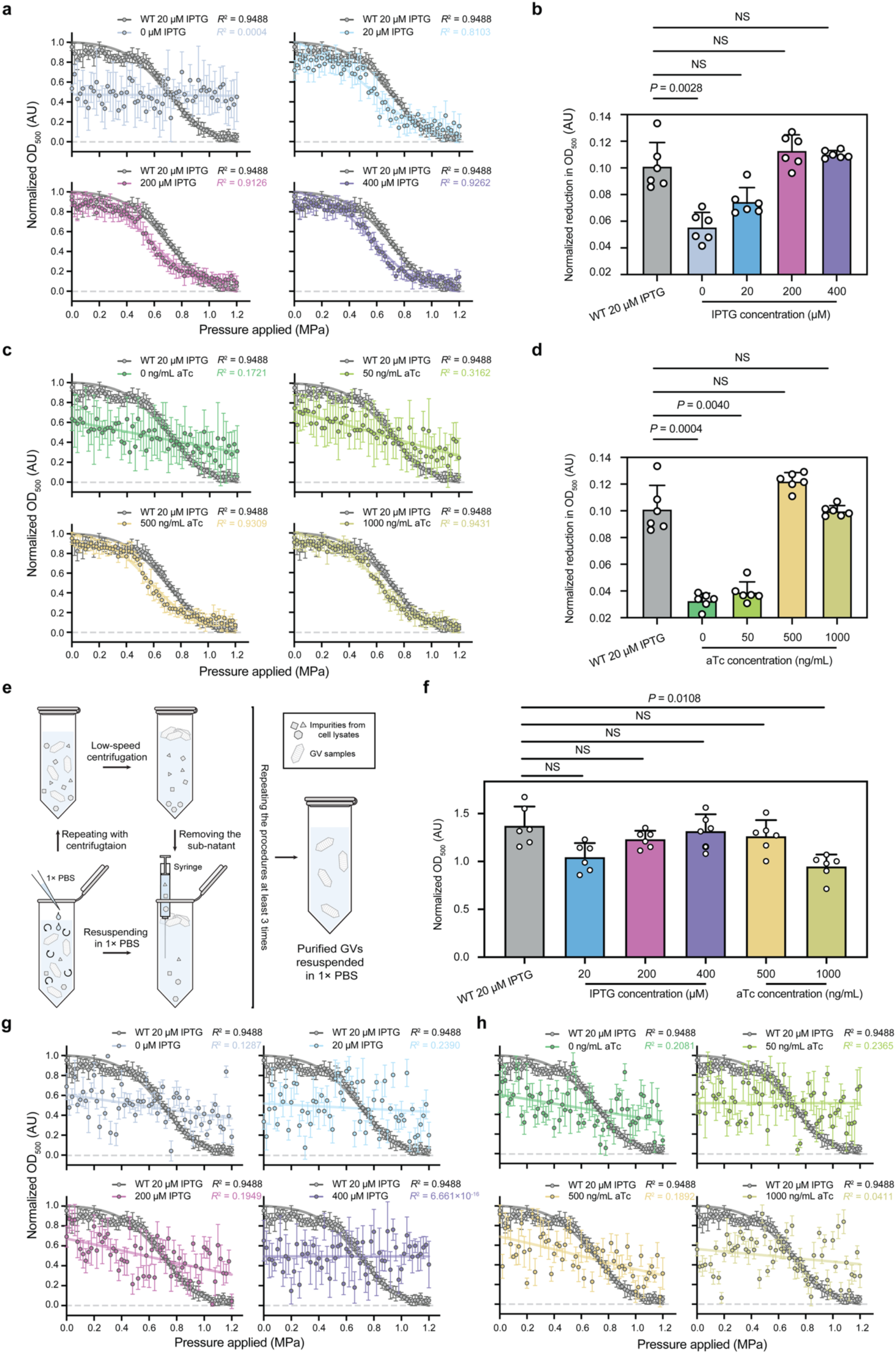
The dual-inducer transcriptional regulation system achieves GV yields comparable to the wild-type operon. **a,** Hydrostatic collapse profiles of *E. coli* cultures expressing GVs under the dual-inducer transcriptional system. Cultures were induced with varying concentrations of IPTG while maintaining aTc at 500 ng/mL, as demonstrated in **Fig. 4a**. WT 20 µM IPTG serves as the control. All data were fitted using a Boltzmann sigmoidal regression model (curves with respective colors), with corresponding *R*² values indicated for each IPTG concentration. **b,** Normalized reduction in OD_500_ values of *E. coli* cultures expressing GVs under the dual-inducer transcriptional regulation system with conditions in **Fig. 4a** and **Fig. 5d**. Statistical analysis was performed against WT 20 µM IPTG using Brown– Forsythe and Welch one-way ANOVA tests, followed by Dunnett T3 multiple comparisons correction. *P* = 0.0028 for 0 µM IPTG (with 500 ng/mL aTc) against WT 20 µM IPTG; NS for all the other groups against WT 20 µM IPTG. **c,** Hydrostatic collapse profiles of *E. coli* cultures expressing GVs under the dual-inducer transcriptional system. Cultures were induced with varying concentrations of aTc while maintaining IPTG at 200 µM, as demonstrated in **Fig. 4c**. WT 20 µM IPTG serves as the control. All data were fitted using a Boltzmann sigmoidal regression model (curves with respective colors), with corresponding *R*² values indicated for each IPTG concentration. **d,** Normalized reduction in OD_500_ values of *E. coli* cultures expressing GVs under the dual-inducer transcriptional regulation system with conditions in **Fig. 4c** and **Fig. 5f**. Statistical analysis was performed against WT 20 µM IPTG using Brown– Forsythe and Welch one-way ANOVA tests, followed by Dunnett T3 multiple comparisons correction. *P* = 0.0004 and *P* = 0.0040 for 0 and 50 ng/mL aTc (with 200 µM IPTG) against WT 20 µM IPTG; NS for all the other groups against WT 20 µM IPTG. **e,** Schematic representation of the buoyancy-based purification procedure for GV samples. The process involves low-speed centrifugation to separate cellular debris, careful removal of the sub-natant, and resuspension of the GV-containing layer in 1× PBS. These steps are repeated in cycles to enhance sample purity. **f,** Normalized OD_500_ values of purified GVs from groups exhibiting high GV yield in **e** and **g**. OD_500_ measurements were normalized to the corresponding *E. coli* culture OD_600_ to account for variations in cell density. Statistical analysis was performed against WT 20 µM IPTG using Brown–Forsythe and Welch one-way ANOVA tests, followed by Dunnett T3 multiple comparisons correction. *P* = 0.0108 for 1000 ng/mL aTc (with 200 µM IPTG) against WT 20 µM IPTG; NS for all the other groups against WT 20 µM IPTG. **g, h** Hydrostatic collapse profiles of *E. coli* cultures expressing GVs under the interchanged dual-inducer transcriptional system (**Sup. Fig. 5**a). (**g**) Cultures were induced with varying concentrations of IPTG while maintaining aTc at 500 ng/mL, as demonstrated in **Sup. Fig. 5**b. (**h**) Hydrostatic collapse profiles of *E. coli* cultures expressing GVs under the interchanged dual-inducer transcriptional system. Cultures were induced with varying concentrations of aTc while maintaining IPTG at 200 µM, as demonstrated in **Sup. Fig. 5**d. WT 20 µM IPTG serves as the control. All data were fitted using a Boltzmann sigmoidal regression model (curves with respective colors), with corresponding *R*² values indicated for each IPTG or aTc concentration. Normalized absorbance values in **a**, **c**, **g**, and **h** are expressed in absorbance units (AU) on the *y-axis*. Data in **a**, **b**, **c**, **d**, **f**, **g**, and **h** represent mean ± s.d for *n* = 6 biologically independent samples for all groups.

We further purified GVs from all experimental groups simultaneously using the centrifugation-assisted flotation^38^ (**Fig. 8e**). After resuspending the purified GVs in equal volumes of 1× PBS, we measured the OD_500_ for each sample and normalized the value to the OD_600_ of the corresponding cultures at harvest. High normalized OD_500_ values indicate greater GVs, as intact GVs effectively scatter light, resulting in increased turbidity^20^. In both the IPTG- and aTc-regulated induction screenings, GVs were successfully purified from all induced groups, except for the non-induced one and the 50 ng/mL aTc group, from which no GVs were recovered in purification. All experimental groups exhibited normalized OD_500_ values comparable to the wild-type control value, although the 1000 ng/mL aTc group displayed marginally reduced normalized OD_500_ values, indicating slightly lower GV production (**Fig. 8f**).

We also assessed the GV yield under the interchanged regulatory configuration (**Fig. 6a**). Employing the previously described screening methodology (**Fig. 6b, d**), we conducted hydrostatic collapse assays on lysed cell cultures. In contrast to the original regulatory system (**Fig. 3b**), none of the experimental groups demonstrated substantial GV production, as indicated by consistently low *R*² values (**Fig. 8g, h**). This outcome is conceivably due to the limited expression of assembly factor proteins under the aTc-inducible system, whose transcriptional strength is insufficient^47^ to express the amount of assembly factors for effective GV assembly. In conclusion, our results demonstrate that the sequential expression of assembly factors prior to shell proteins maintains GV yields comparable to those of the wild-type pNL29 operon, while preserving bacterial viability.

### Time windows of the sequential induction influence cellular stress mitigation and GV yield

Lastly, to investigate the impact of induction timing on the performance of the dual-inducer system in mitigating proteotoxic stress and GV production, we examined whether varying the interval between the initiation of assembly factor protein expression and shell protein expression influences system efficacy. Given that the molar ratio between assembly factors and shell protein is critical for maintaining cellular fitness and proper GV assembly^18, 35, 51^, we screened time intervals between inductions (**Fig. 9a**) to achieve varying amounts of assembly factors and shell protein, and evaluate the influence on mitigating cellular proteotoxic stress and the efficiency of GV formation.

**Fig. 9.**
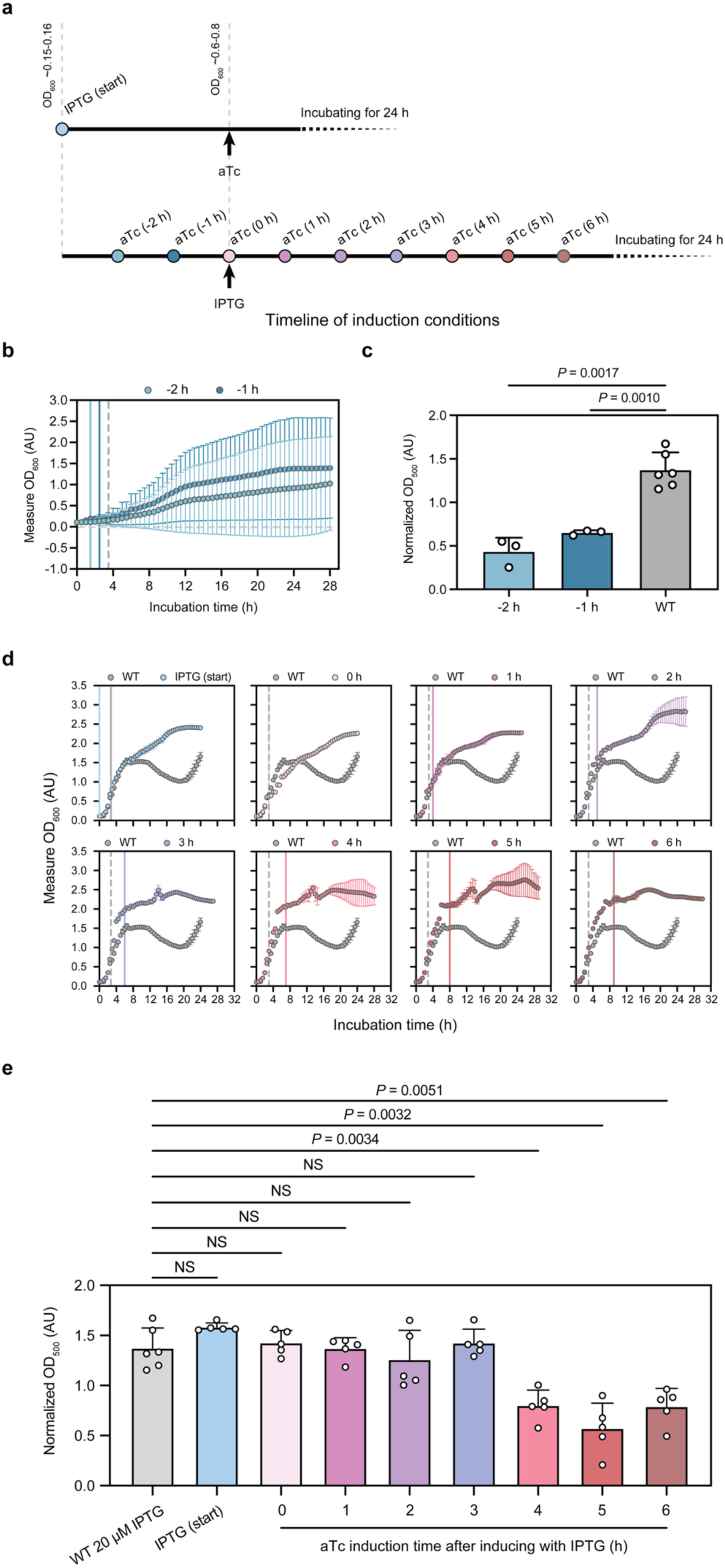
Screening induction timing between assembly factors and shell protein expression reveals an optimal window for maximizing bacterial growth and GV yield. **a,** Schematic timeline illustrating the temporal induction strategies for assembly factor proteins and shell protein expression in *E. coli* using a dual-inducer transcriptional regulation system. In the “IPTG (start)” group (upper), IPTG was present from the initiation of culture, with aTc added upon reaching OD_600_ of 0.6–0.8. The “aTc (−2 h)” and “aTc (−1 h)” groups received aTc 2 hours and 1 hour, respectively, before IPTG addition at OD600 of 0.6–0.8. In the “aTc (0 h)” group, both inducers were added simultaneously at OD_600_ of 0.6–0.8. Subsequent groups, labeled as “aTc (x h),” denote the time intervals (x hours) between IPTG addition at OD_600_ of 0.6–0.8 and subsequent aTc induction. **b,** Measured OD_600_ profiles of *E. coli* cultures expressing GVs under a dual-inducer transcriptional regulation system with various temporal induction strategies. Vertical grey dashed lines indicate the time points at which IPTG was introduced. Vertical colored solid lines denote the time points of aTc addition. Data represent mean ± s.d. for *n* = 5 biologically independent samples for both groups. See **Sup. Fig. 5** for growth curves of individual cultures. **c,** Normalized OD_500_ values of purified GVs from groups exhibiting high GV yield in **b**. OD_500_ measurements were normalized to the corresponding *E. coli* culture OD_600_ to account for variations in cell density. Data represent mean ± s.d. for *n* = 3 biologically independent samples for -2 h and -1 h groups and *n* = 6 biologically independent samples for WT. Statistical analysis was performed against WT using Brown–Forsythe and Welch one-way ANOVA tests, followed by Dunnett T3 multiple comparisons correction. *P* = 0.0017 and 0.0010 for 2- and 1-hour time intervals between aTc addition and subsequent IPTG induction against WT. **d,** Measure OD_600_ profiles of *E. coli* cultures expressing GVs under a dual-inducer transcriptional regulation system with various temporal induction strategies. The single-inducer, wild-type pNL29 GV-expressing *E. coli* culture induced with 20 μM IPTG (WT) served as a control. Vertical grey dashed lines indicate the time points at which IPTG was introduced. Vertical colored solid lines denote the time points of aTc addition. In the “IPTG (start)” group, IPTG was present from the beginning of the culture (vertical blue solid line), and the vertical grey solid line represents the time of aTc addition. Data represent mean ± s.d. for *n* = 5 biologically independent samples for all groups except WT with *n* = 6 biologically independent samples. **e,** Normalized OD_500_ values of purified GVs from groups exhibiting high GV yield in **b**. OD_500_ measurements were normalized to the corresponding *E. coli* culture OD_600_ to account for variations in cell density. Data represent mean ± s.d. for *n* = 5 biologically independent samples except WT 20 µM IPTG with *n* = 6 biologically independent samples. Statistical analysis was performed using Brown–Forsythe and Welch one-way ANOVA tests against WT 20 µM IPTG, followed by Dunnett T3 multiple comparisons correction. *P* = 0.0034, 0.0032, and 0.0051 for 4-, 5-, and 6-hour intervals between IPTG addition and subsequent aTc induction against WT 20 µM IPTG; NS for all the other groups against WT 20 µM IPTG. Absorbance values in **b** and **d** are expressed in absorbance units (AU) on the *y-axis*, representing the amount of light absorbed by the culture at 600 nm wavelength. Normalized absorbance values in **c** are expressed in absorbance units (AU) on the *y-axis*.

In groups where shell protein expression was initiated before assembly factors induction (aTc at -2 h and -1 h; **Fig. 9a**), substantial variability was observed in the growth profiles, as evidenced by significant standard deviations (**Fig. 9b** and **Sup. Fig. 5**). Specifically, two out of five biological replicates in each group exhibited growth arrest due to early GvpA2-induced proteotoxicity. Among the replicates that achieved growth, the purified GV yields were markedly lower than cultures expressing the wild-type pNL29 operon (**Fig. 9c**). These findings suggest that premature expression of GvpA2 without sufficient assembly factor proteins may cause GvpA2 aggregation due to insufficient chaperoning, leading to proteotoxic stress and reduced cell viability. Meanwhile, optical density tracking indicated restored bacterial growth profiles in all remaining groups (**Fig. 9d**). Specifically, the group in which IPTG induction was initiated at the start of culturing exhibited a logarithmic phase (log phase) growth comparable to the wild-type operon-expressing control, yet without experiencing subsequent growth arrest or reductions in optical density (IPTG (start); **Fig. 9d**). This result suggests minimal cellular stress associated with expressing assembly factors alone, and they provided efficient chaperoning of to prevent the proteotoxic stress from expressing GvpA2. The group induced simultaneously for shell and assembly factor proteins displayed a delayed growth rate immediately following induction compared to the wild-type (aTc at 0 h; **Fig. 9d**); however, overall cell viability remained unaffected. This observation could be explained by the comparatively lower transcriptional strength of the aTc-inducible system^47^ relative to the T7-driven system^46, 48^, resulting in lower expression of GvpA2 compared to assembly factors and thus providing sufficient time for assembly factors to handle GvpA2. The highest final optical density values were recorded in the group with assembly factors induction at a 2-hour interval before shell protein induction (aTc at 2 h; **Fig. 9d**). Furthermore, groups with assembly factors induction intervals between 3 and 6 hours exhibited minimal declines in optical density following induction (aTc at 3, 4, 5, and 6 h; **Fig. 9d**). Collectively, these results suggest an optimal induction interval of approximately 2 to 3 hours between assembly factors and shell protein expression to achieve the appropriate stoichiometric balance, effectively mitigate proteotoxic stress, and restore *E. coli* growth. Furthermore, all experimental groups, except those with induction intervals of 4, 5, and 6 hours, demonstrated GV yields comparable to the wild-type control (**Fig. 9e**). The reduced GV yields observed in these longer-interval groups may reflect that cells had already reached high density, which resulted in limited nutrient availability and insufficient resources to support robust shell protein expression, thereby hindering the final yield of GVs. In sum, our results demonstrate the existence of an optimal induction interval between shell assembly factor protein expression, which allows proper stoichiometric balance, mitigates proteotoxic stress, and supports efficient GV assembly. These insights establish a basis for future optimization of GV production in heterologous hosts and may also inform the development of programmable expression systems for other complex macromolecular assemblies.

## Discussion

The protein-based nanostructures, GVs, exhibit unique physical properties that enable them to be repurposed for a range of biomedical applications. Their genetic encodability further allows their use as programmable components within synthetic biology systems. However, the heterologous expression of GVs in microbial hosts encounters challenges due to proteotoxic stress^35, 49^. This stress arises from the aggregation-prone nature of the shell protein GvpA2^40^, which is detrimental to cellular viability. Maintaining robust cellular viability is crucial for biomedical and biotechnological applications. Healthy host cells not only ensure reproducible and reliable results but also directly impact the effectiveness and safety of therapeutic interventions in clinical contexts^52^. In this way, GV-triggered proteotoxic stress compromises the reliability and efficacy of GV-based technologies. To address the proteotoxicity, we developed a dual-inducer transcriptional regulation system engineered to sequentially express assembly factors prior to the induction of GvpA2. This strategy enabled the sufficient accumulation of assembly factors capable of chaperoning GvpA2, facilitating proper folding and GV assembly. As a result, we successfully mitigated proteotoxic effects, restored bacterial growth, and achieved GV yields comparable to those obtained with the wild-type operon.

Our results in interchanged regulatory studies (**Fig. 6** and **Fig. 8g, h**) and temporal induction screenings (**Fig. 9**) indicate that the assembly of GVs relies on precise stoichiometry and temporal relationship among shell and assembly factor proteins in the pNL29 operon. This balance is essential for the successful assembly of GV nanostructures^39^, as an excessively high molar ratio of shell proteins to assembly factors can result in shell protein aggregation, inadequate assembly support, and cytotoxicity, whereas an excessively low ratio may cause inefficient energy utilization due to surplus assembly factors relative to available shell proteins, thus limiting overall GV production.

An important next step is to unravel the underlying molecular mechanism behind this optimal stoichiometry, specifically by investigating the protein complexes and interactions that govern the assembly process. However, studying these relationships is challenging because: (1) the dynamic and transient nature of protein-protein interactions during GV formation^35, 39^ and (2) the difficulty in isolating and quantifying individual proteins within the complex assembly process^35,53^. As understanding these molecule-level interactions is critical for advancing the application of GVs in biotechnology and synthetic biology, recent methodological advances, such as split-GFP assays and affinity-based pull-down techniques^53^, have provided promising approaches to capture transient interactions among GV proteins under physiologically relevant, high-salt conditions. A recent study from our lab advanced this effort by deploying a high-throughput interaction screen in living cells to systematically map interactions among all 11 GV proteins encoded in the pNL29 GV operon^35^. This work resolved key nodes in the protein-protein interaction network, including chaperone-like partnerships and stoichiometric dependencies between structural regulators. These datasets provide a framework for rational engineering of genetic circuits controlling each GV protein in the future, ensuring empirical optimization of cellular physiology, GV yields, and potentially, their physical properties. Engineering GV proteins may also offer a strategy to address challenges in GV assembly. Recent work demonstrated that directed evolution of GvpA in *E. col* can increase GV production and enhanced acoustic signals^54^. In parallel, rational design by leveraging the cryo-EM structure of GvpA^33^ could enable targeted residue modifications to refine solubility, thermostability, or interfacial interactions between assembly components.

Beyond addressing the assembly challenge of GVs, our dual-inducer system may provide a model for optimizing the assembly of other multimeric protein assemblies, ranging from self-assembling protein nanocompartments^55^ to biomolecular condensates^56^ that require finely tuned temporal expression and stoichiometry. For instance, carboxysomes often employ a “core-first” assembly strategy^55^, where sequential incorporation of components is critical for efficient biogenesis. Transferring operons into heterologous hosts may disrupt these established temporal patterns, and synthetic circuits like our dual-inducer system, which enable orthogonal and temporally resolved gene expression, can re-establish native-like sequential logic and promote optimal assembly. Beyond structural systems, this dual-input paradigm is directly applicable to engineered feedback loops and synthetic oscillators. Dual-input regulation has been shown to independently modulate amplitude and period in synthetic gene circuits^57^. Leveraging this capability, our dual-inducer design has the potential to effectively integrate fast and slow feedback motifs^58^ and facilitate robust and noise-resistant switching^59^ behaviors analogous to those observed in natural biological contexts. Therefore, our framework not only facilitates precision in multi-protein assembly but also provides a powerful tool for dynamic regulation in broader synthetic biology applications.

In summary, our study presents a dual-inducer transcriptional regulation system that effectively mitigates proteotoxic stress associated with heterologous expression of GVs in *E. coli* by sequentially controlling assembly factor proteins and the shell protein, GvpA2. This sequential induction restored cellular viability while maintaining GV yields equivalent to wild-type operon expression. The detailed investigation into optimal temporal induction intervals underscored the critical role of stoichiometric balance between shell and assembly factor proteins for efficient GV assembly. These findings establish a tractable framework for further refining synthetic genetic circuits, protein engineering, and mapping the molecular interactions that govern GV biogenesis. By resolving kinetic and stoichiometric bottlenecks, this work advances the potential for tunable and scalable production of GV-based tools for biomedical engineering and synthetic biology.

## Methods

### Plasmids construction

The plasmid pST39-pNL29, encoding the gas vesicle (GV) gene cluster from *Priestia megaterium* (formerly *Bacillus megaterium*), was obtained from Addgene (plasmid #91696). The GV gene cluster sequence corresponds to GenBank accession number AF053765.1. The major shell protein, GvpA2, is derived from *P. megaterium* and is cataloged under UniProt ID O68677 (GVPA2_PRIMG). Structural representations of GvpA2, as depicted in **Fig. 2a, b** by using ChimeraX (version 1.9), were adapted from cryo-electron microscopy (cryo-EM) data available in the Protein Data Bank (PDB 7R1C)^33^. The plasmid pAJM.011, incorporating the Tet repressor (TetR) and the tetracycline-responsive promoter (aTc-inducible system)^60^, was sourced from Addgene (plasmid #108529). The monomeric Cherry red fluorescent protein (mCherry) gene was obtained from Addgene (plasmid #29747), and the superfolder green fluorescent protein (sfGFP) gene was acquired from Addgene (plasmid #85492). Oligonucleotides for molecular cloning were synthesized by Integrated DNA Technologies (IDT, Coralville, IA). All genes of interest were cloned into the pST39 vector backbone using the NEBuilder® HiFi DNA Assembly Master Mix (New England Biolabs, Ipswich, MA), following the manufacturer’s protocol.

### Recombinant GV expression in *E. coli*

Chemically competent BL21 Star™ (DE3)pLysS One Shot™ *E. coli* strain (Thermo Fisher Scientific, Waltham, MA) was transformed with 1 µL of 10 ng/µL plasmid DNA—either wild-type pST39-pNL29 and GvpA2 only or dual-inducer transcriptional regulation system variants—using the heat shock method. The plasmid DNA was gently mixed with 10 µL of competent cells and incubated on ice for 30 minutes. Cells were then subjected to a 30-second heat shock at 42 °C, followed by immediate placement on ice for 5 minutes. Subsequently, 400 µL of pre-warmed SOC medium (Thermo Fisher Scientific, Waltham, MA) was added to each transformation reaction, and the cells were incubated at 37 °C for 1 hour with shaking at 250 rpm to allow recovery. Transformed cells were then plated on LB (Luria-Bertani) Miller agar plates (Thermo Fisher Scientific, Waltham, MA) with 1% glucose, 25 µg/mL chloramphenicol (MilliporeSigma, Burlington, MA), and 100 µg/mL carbenicillin (Gold Biotechnology, Olivette, MO). Plates were incubated overnight at 30 °C. Single colonies were selected and cultured in 3 mL of LB Miller broth containing the same concentrations of glucose and antibiotics in 24-deep-well plates (Thermo Fisher Scientific, Waltham, MA) at 30 °C for 16 hours with shaking at 600 rpm on a microplate incubator (Incu-mixer™ MP Heated Microplate Vortexer; Benchmark Scientific Inc, Sayreville, NJ). For GV expression, overnight cultures were diluted to an initial optical density at 600 nm (OD_600_) of 0.05 in 3 mL of 2×YT (2× Yeast Extract Tryptone medium) broth (Thermo Fisher Scientific, Waltham, MA) within 24-deep-well plates. All cell cultures were incubated at 30°C with shaking at 600 rpm until an optical density (OD_600)_ of 0.6–0.8 was reached. For cells with the wild-type pNL29 plasmid, isopropyl β-D-1-thiogalactopyranoside (IPTG; Teknova, Hollister, CA) was added to final concentrations of 0, 20, 200, or 400 µM to induce expression. For cells with dual-inducer transcriptional regulation system variants plasmids, induction was performed with either IPTG or anhydrotetracycline (aTc; MilliporeSigma, Burlington, MA) with concentrations specified in the figure legends following the same OD_600_ threshold. After an additional 3 hours of incubation under the same conditions, a second inducer (IPTG or aTc) was added as per experimental design.

150 µL of culture was transferred into wells of the 96-well black, flat-bottom polystyrene microplate (Corning® 96-well Black Flat Bottom Polystyrene Not Treated Microplate 3631; Corning Inc., Corning, NY) for each measurement. Cell growth was tracked by measuring OD_600_ using the TECAN Spark multimode microplate reader (Tecan Group Ltd., Männedorf, Switzerland).

### Isolation, purification, and quantification of GVs

*E. coli* cultures expressing GVs were harvested and subjected to low-speed centrifugation to exploit the buoyant properties of GVs. Cultures were centrifuged at 400 × *g* for 4 hours at 4 °C in 2.0 mL Eppendorf™ Safe-Lock tubes (Thermo Fisher Scientific, Waltham, MA), with a sample volume of 1.5 mL per tube. The intermediate layer between the buoyant GV-containing fraction and the sedimented cells was carefully removed using a syringe. The remaining cell pellet was resuspended in SoluLyse-Tris buffer (Genlantis, San Diego, CA) at a volume of 60 µL per 1.5 mL of initial culture. Lysozyme (MilliporeSigma, Burlington, MA) and RNase-free Deoxyribonuclease I (DNase I; MilliporeSigma, Burlington, MA) were added to final concentrations of 250 µg/mL and 10 µg/mL, respectively, to facilitate cell lysis and nucleic acid degradation. The suspension was incubated with gentle rotation at 4 °C for 30 minutes. Post-lysis, the samples underwent a second centrifugation at 400 × *g* for 2 hours at 4 °C to separate the buoyant GVs from cellular debris, as depicted in **Fig. 8e**. The buoyant GVs accumulated at the top of the liquid surface were carefully collected by aspirating the underlying supernatant and cell debris using a syringe. The isolated GVs were subjected to three successive washes with 1× PBS (Teknova, Hollister, CA), each followed by centrifugation under the same conditions.

Purified GV samples were standardized to a final volume of 1.5 mL by resuspension in 1× PBS. The OD_500_ was measured using a Take3 microvolume plate on an Agilent BioTek Synergy H4 Hybrid microplate reader (Agilent Technologies, Santa Clara, CA). The measured OD_500_ values were normalized by dividing by the corresponding OD_600_ of the *E. coli* cultures before lysis to account for variations in cell density at the time of harvest.

### Fluorescent reporter protein expression in *E. coli*

Plasmids encoding either dual-inducer transcriptional regulation system variants or single-reporter constructs were transformed into chemically competent *E. coli* cells using standard heat-shock protocols, as mentioned in the previous section. Following transformation, cells were incubated and cultured under the conditions previously described. Upon reaching OD_600_ of 0.6–0.8, induction strategies were applied according to the experimental designs outlined in **Fig. 3** and **Sup. Figs. 2–4**.

Fluorescence intensity measurements of *E. coli* cultures expressing fluorescent reporter proteins were conducted using a multimode microplate reader. A volume of 150 µL from each culture was transferred into individual wells of a 96-well black, flat-bottom polystyrene microplate (Corning® 96-well Black Flat Bottom Polystyrene Not Treated Microplate 3631; Corning Inc., Corning, NY). For mCherry fluorescence detection, the excitation wavelength was set at 587 nm, and the emission was detected at 610 nm, with a bandwidth of 5 nm. For sfGFP fluorescence detection, the excitation wavelength was 488 nm, and the emission wavelength was 510 nm, with a bandwidth of 5 nm. Gain settings were optimized to 150 for both to ensure signal linearity and prevent detector saturation. All measurements were performed at room temperature, and data acquisition was managed using the SparkControl software (version 3.1 SP1). Fold change in fluorescence intensity values in **Fig. 3** and **Sup. Figs. 2–4** were calculated using the equation: Fold change in fluorescence intensity = [(measured fluorescence intensity of studied cells) / (OD_600_ of culture at harvest of studied cells)] / [(measured fluorescence intensity of not induced cells) / (OD_600_ of culture at harvest of not induced cells)].

### Temporal induction of assembly factors protein and shell protein in *E. coli*

Transformation of plasmids into chemically competent *E. coli* cells, subsequent incubation, and GV expression were performed following the protocols and conditions described in previous sections. Various induction strategies were implemented based on the experimental design. In the “IPTG (start)” condition (**Fig. 9**), IPTG (Teknova, Hollister, CA) was added at the initiation of culture, and aTc (MilliporeSigma, Burlington, MA) was introduced when cultures reached OD_600_ of 0.6–0.8. For the “aTc (−2 h)” and “aTc (−1 h)” conditions (**Fig. 9**), aTc was administered 2 hours and 1 hour, respectively, before IPTG induction at OD_600_ of 0.6–0.8. In the “aTc (0 h)” condition (**Fig. 9**), both IPTG and aTc were simultaneously added upon cultures reaching OD_600_ of 0.6–0.8. Additional induction schemes (**Fig. 9**), labeled as “aTc (x h),” involved the addition of aTc at defined time intervals (x hours) following IPTG induction at OD_600_ of 0.6–0.8. Following induction, GVs were purified from each experimental group and quantified using the previously described protocols.

### Transmission electron microscopy

*E. coli* cell samples were diluted to OD_600_ of 0.1 in 1× PBS. A 5 µL aliquot of the diluted suspension was applied to a 200-mesh carbon-coated copper grid (Ted Pella) and incubated for 3 minutes to allow adherence. Excess liquid was gently removed by blotting with Whatman filter paper (MilliporeSigma, Burlington, MA). For negative staining of purified GVs, samples were diluted to OD_500_ of 0.2 in 1× PBS. A 5 µL aliquot of the diluted sample was applied to a 200-mesh carbon-coated copper grid (Ted Pella, Redding, CA) and incubated for 3 minutes. Excess liquid was carefully blotted away using Whatman filter paper (MilliporeSigma, Burlington, MA). Subsequently, the grids were stained with a 5 µL drop of 2% (w/v) uranyl acetate solution (Electron Microscopy Sciences, Hatfield, PA) for contrast enhancement. After staining, excess stain was removed by blotting with filter paper. The grids were then air-dried in a chemical fume hood with lateral ventilation for 2 hours to ensure complete drying. High-resolution TEM imaging was performed using a JEOL JEM-2100 Field Emission Gun TEM and an FEI Tecnai F20 TEM.

### Hydrostatic collapse assay

The experimental setup for hydrostatic collapse assay with pressurized absorbance spectroscopy was configured as previously described^38^. The system comprised (**Fig. 7a**): (1) a computer running MATLAB scripts to control pressure application; (2) a compressed nitrogen gas cylinder equipped with control valves and a pressure regulator; (3) a single-valve pressure controller (PC series; Alicat Scientific, Tucson, AZ) interfaced with the computer; (4) a quartz cuvette (176.700-QS; Hellma GmbH & Co. KG, Müllheim, Germany); and (5) a UV–Vis spectrophotometer (STS-VIS; Ocean Optics, Orlando, FL) with an integrated light source and cuvette holder. Purified GVs were diluted to OD_500_ of 0.2 in 1× PBS. *E. coli* cells expressing GVs and other GV variants were lysed following the protocol detailed in the previous section and treated with 6 M urea. Samples were loaded into a pressurizable cuvette. OD_500_ was measured using the spectrophotometer, while the cuvette was subjected to increasing pressure via a single-valve pressure controller. The pressure was incrementally increased from 0 to 1.4 MPa in 0.2 MPa steps, allowing for the assessment of GV collapse under controlled hydrostatic pressure conditions.

OD_500_ were normalized using the following equation: Normalized OD_500_ = (Measured OD_500_ - Minimum OD_500_) / (Maximum OD_500_ - Minimum OD_500_). Hydrostatic collapse profiles were fitted using a Boltzmann sigmoidal function, with constraints set such that the bottom equals 0 and the top equals 1, to model the transition from intact to collapsed states. A coefficient of determination (*R*²) of this nonlinear fit was provided. To quantify GV expression levels in *E. coli* while accounting for variations in cell density, we normalized the reduction in OD500, calculated as follows: Normalized reduction in OD_500_ = (Maximum OD_500_ - Minimum OD_500_) / OD_600_ at harvest. This metric provides a standardized measure of GV production per unit of cellular biomass.

### Live-cell fluorescence and phase contrast microscopy

*E. coli* cells expressing mCherry, mCherry::GvpA2, or GvpA2::mCherry were grown under conditions previously described. 4 hours post-induction with IPTG cells were pelleted by centrifugation and washed three times with Hank’s Balanced Salt Solution (HBSS; Thermo Fisher Scientific, Waltham, MA). The cell suspension was then diluted in HBSS and applied to 35 mm glass-bottom dishes with number 1.5 coverslips (MatTek Life Sciences, Ashland, MA) for imaging. Live-cell fluorescence imaging was performed using a ZEISS Elyra 7 microscope (Carl Zeiss AG, Germany) equipped with SIM² Lattice Structured Illumination Microscopy (SIM) and Total Internal Reflection Fluorescence (TIRF) capabilities. The system was configured with a Plan-Apochromat 63×/1.46 oil immersion objective lens. mCherry fluorescence signal was excited using a 561 nm laser line, and emission was collected between 580–620 nm using a PCO.edge sCMOS camera. Image acquisition and processing were performed using ZEISS ZEN Blue (version 3.11) and Fiji (ImageJ2).

*In vitro* protein synthesis reactions were analyzed using phase contrast microscopy to assess sample morphology. A 5 μL aliquot of each reaction mixture was applied to a 35 mm glass-bottom dish with a No. 1.5 coverslip (MatTek Life Sciences, Ashland, MA). Imaging was acquired on a Nikon Eclipse Ti2 microscope (Nikon, Japan) equipped with a 40×/0.60 NA objective lens and a phase contrast ring corresponding to Ph2. The microscope was configured for Köhler illumination, and phase contrast images were captured using a PCO.edge sCMOS camera. Image acquisition and processing were conducted using Nikon NIS-Elements software (version 6.10. 01) and Fiji (ImageJ2).

### Quantification and statistical analysis

The figures and figure captions provide details regarding biologically independent sample size (*n*), *P*-values, and other statistical information. All statistical analyses were conducted using GraphPad Prism software (version 10.4.2). Data are presented as mean ± standard deviation (s.d.). Comparisons between two groups were evaluated using a two-tailed unpaired Student’s *t*-test with Welch’s correction to account for unequal variances. For multiple group comparisons, the Brown– Forsythe and Welch one-way analysis of variance (ANOVA) with Dunnett’s T3 correction was employed, which does not assume equal variance across groups. Statistical significance was defined as *P* < 0.05.

### Statistics and reproducibility

All representative micrographs are based on experiments repeated independently at least three times (*n* = 3 biologically independent samples).

## Acknowledgments

We thank the Shared Equipment Authority (SEA) at Rice University for access to core facilities and instrumentation, and Dr. Wenhua Guo, Dr. Alloysius Budi Utama, and Lihua Ma for technical training. This work was supported by the Cancer Prevention and Research Institute of Texas (CPRIT, RR190081), the National Institutes of Health (NIH, R35GM155015 and R21EB033607), the Welch Foundation (C-2249 and C-2069), the G. Harold and Leila Y. Mathers Foundation, the John S. Dunn Foundation, and the Open Collective Foundation.

## Author information

## Authors and Affiliations

**Department of Bioengineering, Rice University, Houston, TX 77005, USA**

Zongru Li, Sumin Jeong, and George J. Lu

## Contributions

Conceptualization, G.J.L.; Methodology, G.J.L., Z.L., S.J.; Investigation, Z.L., S.J.; Formal Analysis, Z.L., S.J.; Manuscript writing, G.J.L., Z.L., S.J.; Supervision and Funding Acquisition, G.J.L. All authors have reviewed and approved the final version of the manuscript.

## Ethics declarations

All authors meet the authorship criteria as defined by Nature Portfolio journals, having made substantial contributions to the conception, design, execution, or interpretation of the study. Roles and responsibilities were agreed upon prior to the commencement of the research. Contributors who did not meet all criteria for authorship are acknowledged in the Acknowledgements section. This research was conducted in collaboration with local partners to ensure relevance and applicability to the local context. The study design and implementation were carried out with respect for local norms and practices. Efforts were made to prevent any form of stigmatization, discrimination, or personal risk to participants. Relevant local and regional research has been appropriately cited to acknowledge existing contributions and to provide context. Relevant local and regional research has been appropriately cited to acknowledge existing contributions and to provide context.

---

**Supplementary Figure 1.**
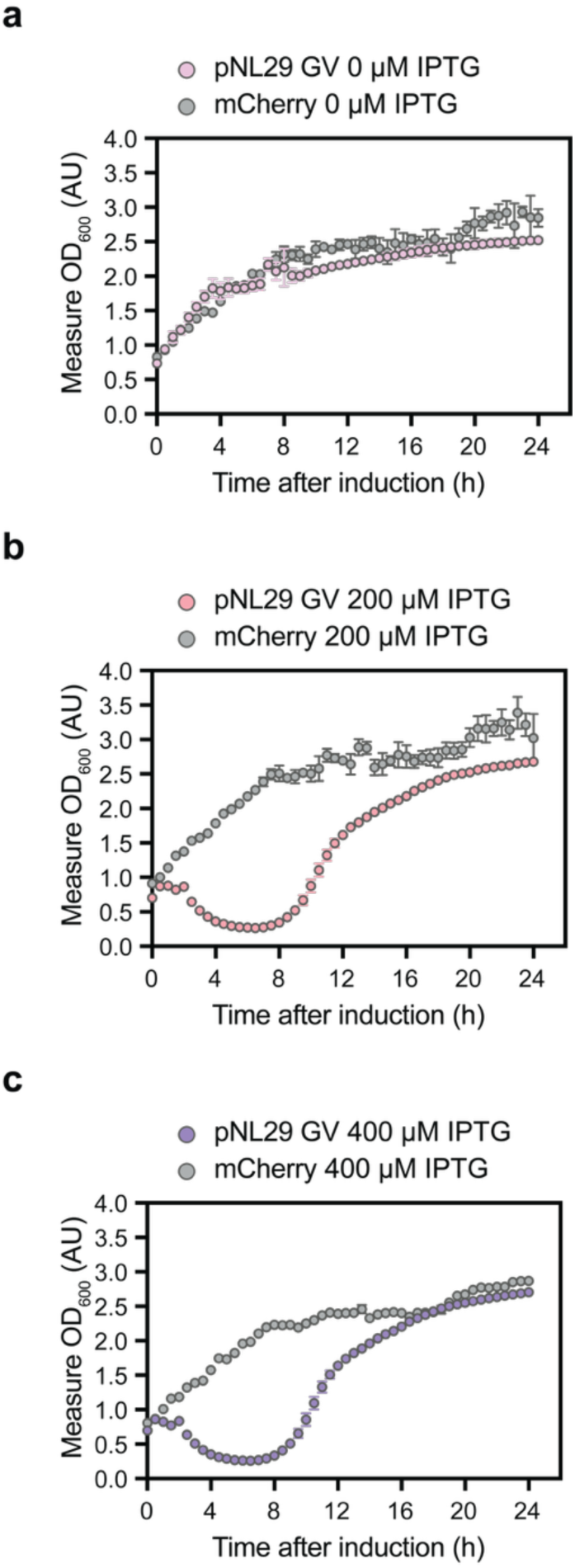
Growth profile of GV-expressing *E. coli* under varying IPTG induction concentrations. **a, b, c,** Growth of *E. coli* cultures after being induced with 0 µM (**a**), 200 µM (**b**), and 400 µM (**c**) IPTG was monitored by measuring OD_600_. Data represent mean ± standard deviation (s.d.) for *n* = 6 biologically independent samples for the pNL29 GV-expressing culture and *n* = 4 biologically independent samples for the mCherry-expressing culture. Absorbance values in **a**, **b,** and **c** are expressed in absorbance units (AU) on the *y-axis*, representing the amount of light absorbed by the culture at 600 nm wavelength.

**Supplementary Figure 2.**
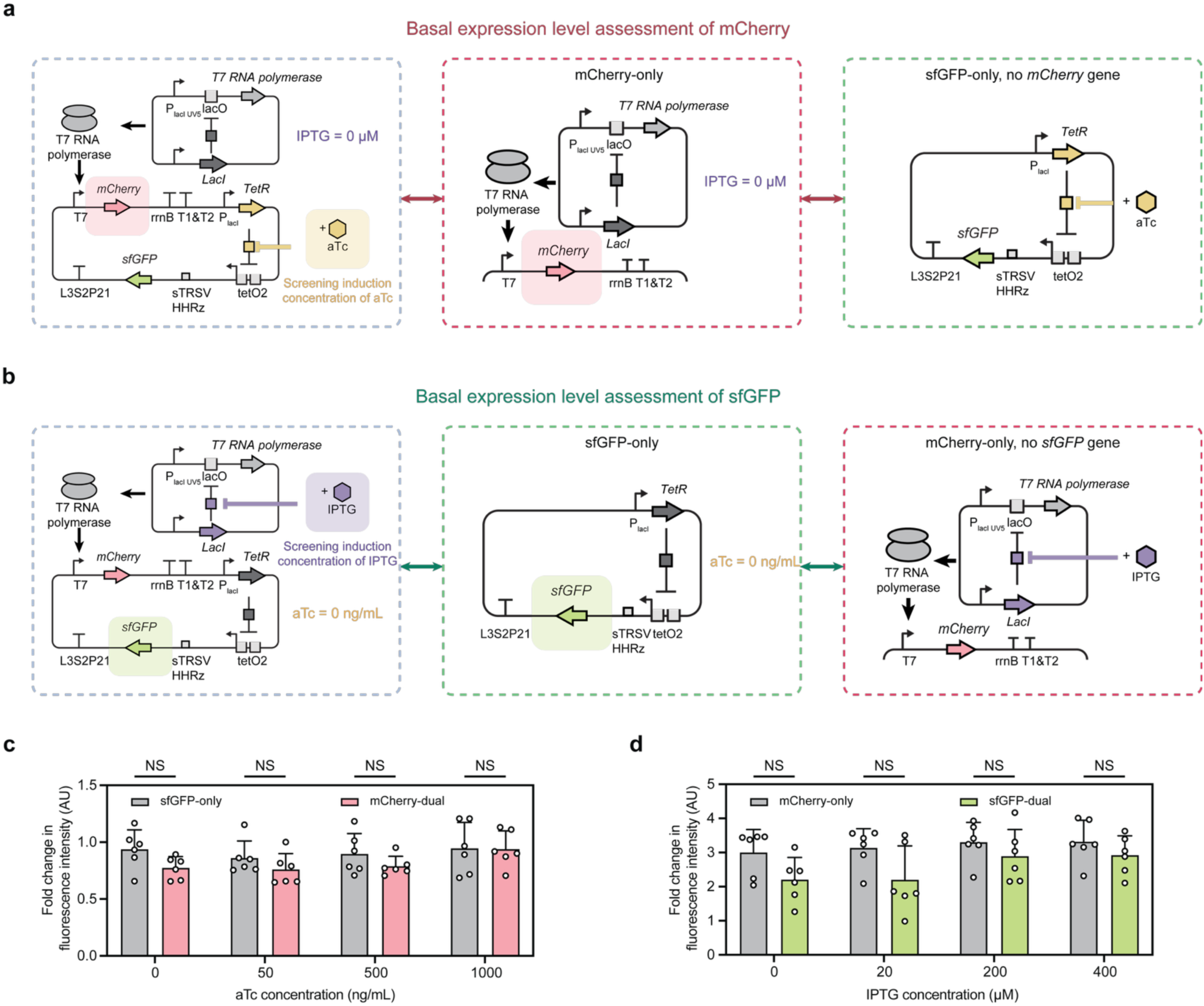
Basal expression levels of individual regulatory elements within the dual-inducer transcriptional regulation system were assessed using fluorescent reporters. **a, b,** Basal expression levels of individual regulatory elements within the dual-inducer transcriptional regulation system were assessed using fluorescent reporters and compared to single-reporter constructs. (**a**) The basal expression of mCherry was evaluated under 0 µM IPTG with varying aTc concentrations (blue dashed box). Fold change in fluorescence intensity was compared to single-reporter constructs: mCherry-only induced with 0 µM IPTG (red dashed box; data in **Fig. 3f**) and sfGFP-only induced with varying aTc concentrations (green dashed box; data in **c**). (**b**)The basal expression of sfGFP was evaluated under 0 ng/mL aTc with varying IPTG concentrations (blue dashed box). Fold change in fluorescence intensity was compared to single-reporter constructs: sfGFP-only induced with 0 ng/mL aTc (green dashed box; data in **Fig. 3e**) and mCherry-only induced with varying IPTG concentrations (green dashed box; data in **d**). **c, d,** Fold change in fluorescence intensity comparing the dual-inducer transcriptional regulation system to single-reporter constructs lacking the corresponding reporter gene, to assess basal expression levels. **(c)** mCherry-dual groups (0 µM IPTG) were compared to sfGFP-only groups across increasing aTc concentrations (0, 50, 500, and 1000 ng/mL). **(d)** sfGFP-dual groups (0 ng/mL aTc) were compared to mCherry-only groups across increasing IPTG concentrations (0, 20, 200, and 400 µM). Fold change in fluorescence intensity values in **c** and **d** are expressed in arbitrary units (AU) on the *y-axis*. Data in **c** and **d** represent mean ± s.d. for *n* = 6 biologically independent samples for all groups. The statistical significance between groups in **c** and **d** was tested using a two-tailed unpaired Student’s *t*-test with Welch correction. NS for all comparisons in **c** and **d**.

**Supplementary Figure 3.**
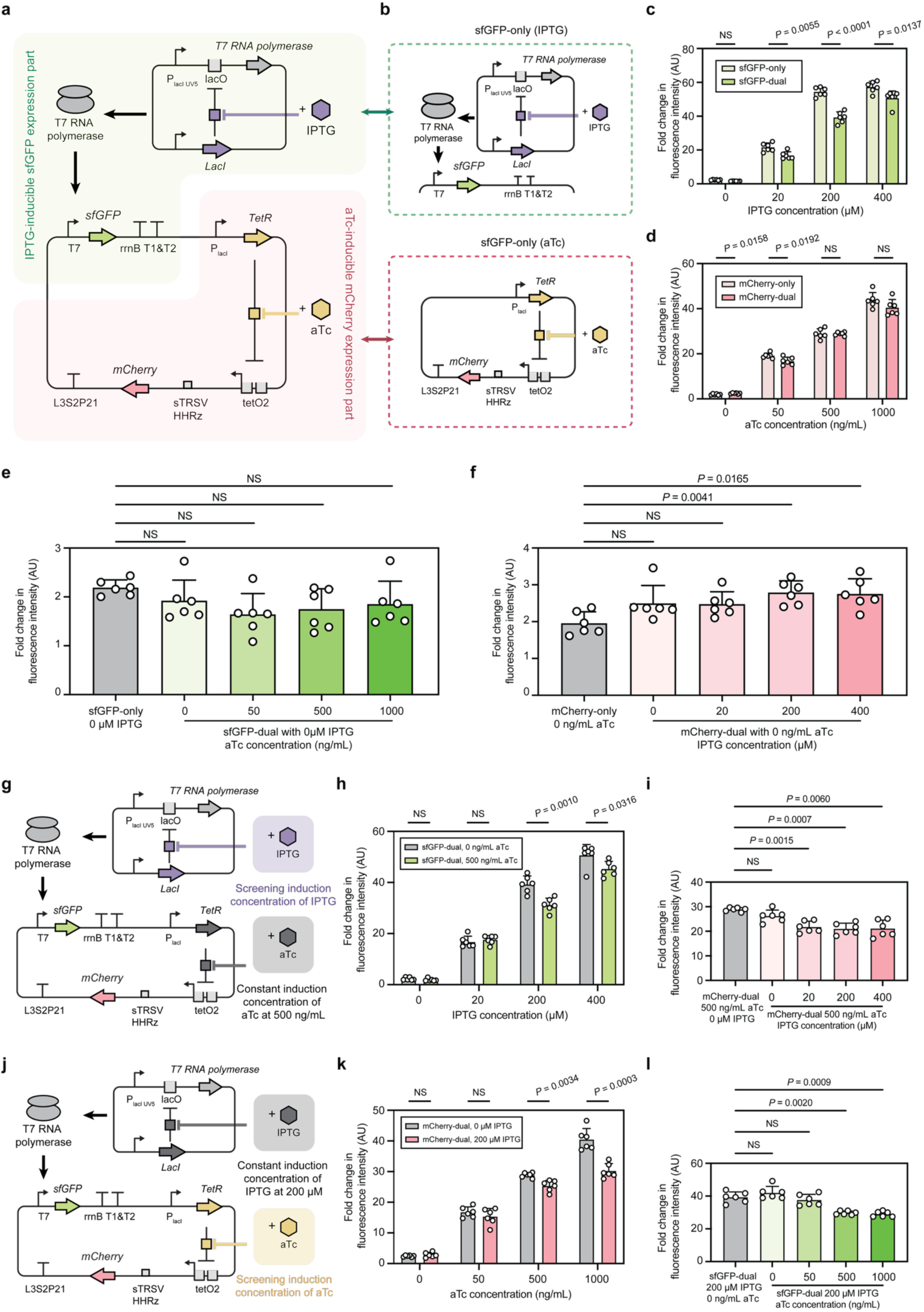
Interchanging regulatory control of fluorescent reporters in the dual-inducer transcriptional regulation system resulted in expression patterns comparable to the original configuration. **a, b,** Schematic representation of transcriptional regulation systems for expression of fluorescent reporter proteins. (**a**) The interchanged dual-inducer transcriptional regulation system enables separate control of sfGFP and mCherry expression. The sfGFP expression is driven by a T7 promoter and activated upon IPTG induction (green background); mCherry expression is regulated by an aTc-inducible system, repressed by TetR in the absence of aTc and derepressed upon aTc addition (red background). (**b**) Two single-reporter genetic constructs, with sfGFP expressed upon IPTG induction (“sfGFP-only (IPTG),” green dashed box) and mCherry expressed upon aTc induction (“mCherry-only (aTc),” red dashed box), respectively. **c, d,** Fold change in fluorescence intensity comparing the dual-inducer transcriptional regulation system to single-reporter constructs (**b**) under varying inducer concentrations. In the dual system, only the inducer corresponding to the respective reporter was added, resulting in “sfGFP-dual” (**c**) and “mCherry-dual” (**d**). NS for sfGFP-only vs. sfGFP-dual at 0 µM IPTG; *P* = 0.0055, < 0.0001, and = 0.0137 at 20, 200, and 400 µM IPTG, respectively. *P* = 0.0158 and 0.0137 for mCherry-only vs. mCherry-dual at 0 and 50 ng/mL aTc. NS for comparisons at 500 and 1000 ng/mL aTc. **e, f,** Fold change in fluorescence intensity comparing the dual-inducer transcriptional regulation system to single-reporter constructs to assess the basal expression level. (**e**) sfGFP-dual groups (0 μM IPTG) with varying aTc concentrations (0, 50, 500, 1000 ng/mL) were compared to sfGFP-only at 0 μM IPTG. (**f**) mCherry-dual groups (0 ng/mL aTc) with varying IPTG concentrations (0, 20, 200, 400 μM) were compared to mCherry-only at 0 ng/mL aTc. For sfGFP-dual, NS for all comparisons. For mCherry-dual, NS for comparisons at 0 and 20 µM IPTG; *P* = 0.0041 and 0.0165 at 200 and 400 μM IPTG. **g,** Schematic representation of the genetic construct for fluorescent reporter proteins under the dual-inducer transcriptional regulation system. Cultures were induced with varying concentrations of IPTG (0, 20, 200, and 400 μM) while maintaining aTc at a constant concentration of 500 ng/mL. **h,** Fold change in sfGFP fluorescence intensity in the dual-inducer transcriptional system with and without aTc. sfGFP-dual groups with 0 ng/mL aTc were compared to groups with 500 ng/mL aTc across varying IPTG concentrations (**g**). NS for comparisons at 0 and 20 µM IPTG; *P* = 0.0010 and 0.0316 for comparisons at 200 and 400 µM IPTG. **i,** Fold change in mCherry fluorescence intensity in the dual-inducer transcriptional system with and without IPTG. mCherry-dual groups with 500 ng/mL aTc and 0 µM IPTG were compared to those with 500 ng/mL aTc across varying IPTG concentrations. NS for comparisons at 0 µM IPTG; *P* = 0.0015, 0.0007, and 0.0060 for comparisons at 20, 200, and 400 µM IPTG. **j,** Schematic representation of the genetic construct for fluorescent reporter proteins under the dual-inducer transcriptional regulation system. Cultures were induced with varying concentrations of aTc (0, 50, 500, and 1000 ng/mL) while maintaining IPTG at a constant concentration of 200 µM. **k,** Fold change in mCherry fluorescence intensity in the dual-inducer transcriptional system with and without aTc. mCherry-dual groups with 0 µM IPTG were compared to groups with 200 µM IPTG across varying aTc concentrations (**j**). NS for comparisons at 0 and 50 ng/mL aTc; *P* = 0.0034 and 0.0003 for comparisons at 500 and 1000 ng/mL aTc. **l,** Fold change in sfGFP fluorescence intensity in the dual-inducer transcriptional system with and without IPTG. sfGFP-dual groups with 200 µM IPTG and 0 ng/mL aTc were compared to those with 200 µM IPTG across varying aTc concentrations. NS for comparisons at 0 and 50 ng/mL aTc; *P* = 0.0020 and 0.0009 for comparisons at 500 and 1000 ng/mL aTc. Fold change in fluorescence intensity values in **c**, **d, e**, **f, h**, **i, k,** and **l** are expressed in arbitrary units (AU) on the *y-axis*. Data in **c**, **d, e**, **f, h**, **i, k,** and **l** represent mean ± s.d. for *n* = 6 biologically independent samples for all groups. The statistical significance between groups in **c**, **d, h**, and **k** was tested using a two-tailed unpaired Student’s *t*-test with Welch correction. The statistical significance was tested against control groups (grey) in **e**, **f**, **i,** and **l** using Brown–Forsythe and Welch one-way ANOVA tests, followed by Dunnett T3 multiple comparisons correction. *P* values of each comparison are indicated in each caption.

**Supplementary Figure 4.**
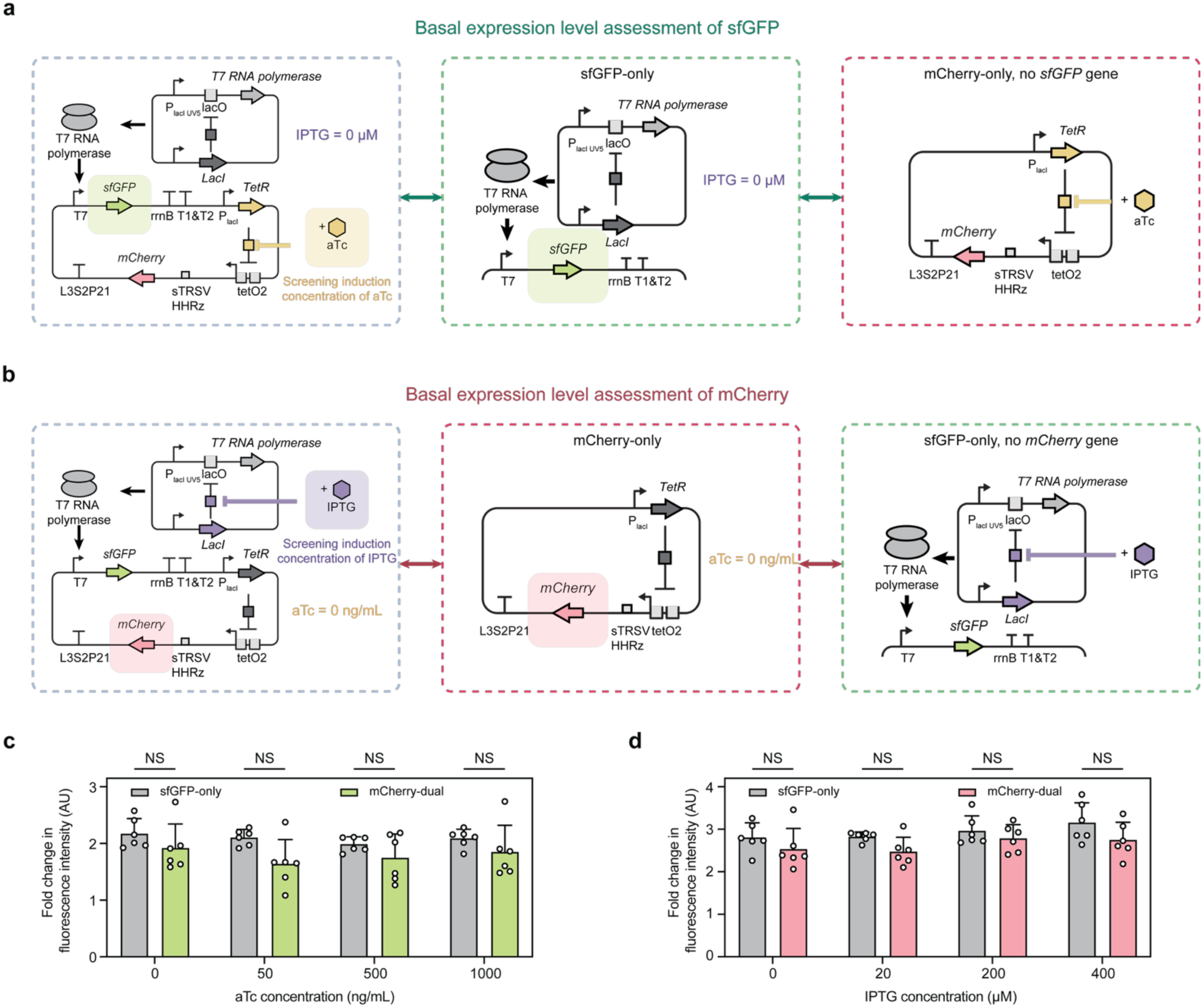
Basal expression levels of individual regulatory elements within the dual-inducer transcriptional regulation system were assessed using interchanged fluorescent reporters. **a, b,** Basal expression levels of individual regulatory elements within the interchanged dual-inducer transcriptional regulation system were assessed using fluorescent reporters and compared to single-reporter constructs. (**a**) The basal expression of sfGFP was evaluated under 0 µM IPTG with varying aTc concentrations (blue dashed box). Fold change in fluorescence intensity was compared to single-reporter constructs: sfGFP-only induced with 0 µM IPTG (green dashed box; data in **Sup. Fig. 3**f) and mCherry-only induced with varying aTc concentrations (red dashed box; data in **c**). (**b**)The basal expression of mCherry was evaluated under 0 ng/mL aTc with varying IPTG concentrations (blue dashed box). Fold change in fluorescence intensity was compared to single-reporter constructs: mCherry-only induced with 0 ng/mL aTc (red dashed box; data in **Sup. Fig. 3**e) and sfGFP-only induced with varying IPTG concentrations (green dashed box; data in **d**). **c, d,** Fold change in fluorescence intensity comparing the interchanged dual-inducer transcriptional regulation system to single-reporter constructs lacking the corresponding reporter gene, to assess basal expression levels. **(c)** sfGFP-dual groups (0 µM IPTG) were compared to mCherry-only groups across increasing aTc concentrations (0, 50, 500, and 1000 ng/mL). **(d)** mCherry-dual groups (0 ng/mL aTc) were compared to sfGFP-only groups across increasing IPTG concentrations (0, 20, 200, and 400 µM). Fold change in fluorescence intensity values in **c** and **d** are expressed in arbitrary units (AU) on the *y-axis*. Data in **c** and **d** represent mean ± s.d. for *n* = 6 biologically independent samples for all groups. The statistical significance between groups in **c** and **d** was tested using a two-tailed unpaired Student’s *t*-test with Welch correction. NS for all comparisons in **c** and **d**.

**Supplementary Figure 5.**
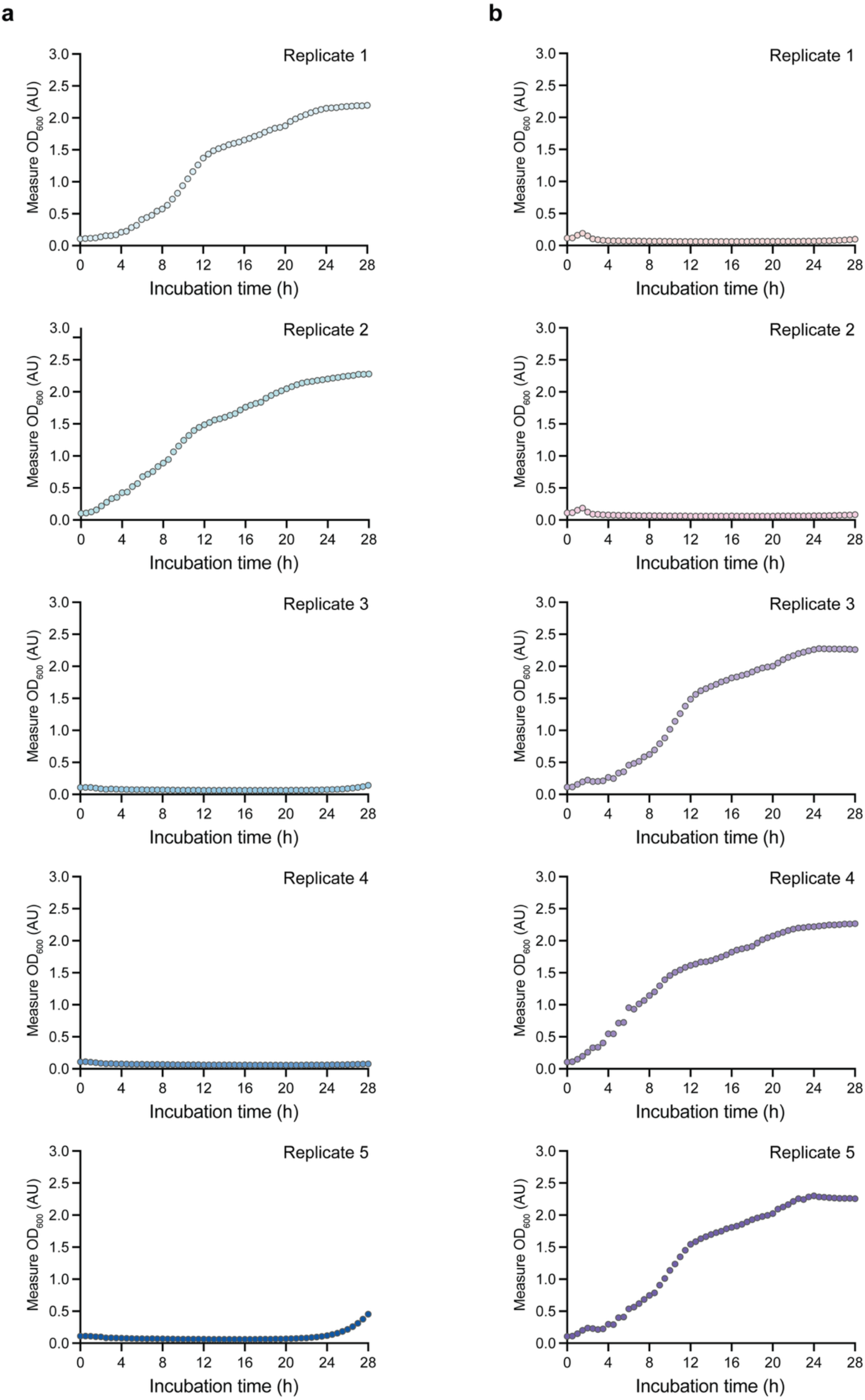
Growth profile of GV-expressing *E. coli* under a dual-inducer transcriptional regulation system with shell protein induced before assembly factors. **a, b,** Measure OD_600_ profiles of *E. coli* cultures expressing GVs under a dual-inducer transcriptional regulation system with various temporal induction strategies, in which groups received aTc 2 hours (**a**) and 1 hour (**b**), respectively, before IPTG addition at OD600 of 0.6–0.8. Each plot represents OD_600_ values for one biologically independent sample. Absorbance values in **a** and **b** are expressed in absorbance units (AU) on the *y-axis*, representing the amount of light absorbed by the culture at 600 nm wavelength. Normalized absorbance values in **c** are expressed in absorbance units (AU) on the *y-axis*.

